# Solvent quality and chromosome folding in *Escherichia coli*

**DOI:** 10.1101/2020.07.09.195560

**Authors:** Yingjie Xiang, Ivan V. Surovtsev, Yunjie Chang, Sander K. Govers, Bradley R. Parry, Jun Liu, Christine Jacobs-Wagner

## Abstract

All cells must fold their genomes, including bacterial cells where the chromosome is compacted into a domain-organized meshwork called nucleoid. Polymer conformation depends highly on the quality of the solvent. Yet, the solvent quality for the DNA polymer inside cells remains unexplored. Here, we developed a method to assess this fundamental physicochemical property in live bacteria. By determining the DNA concentration and apparent average mesh size of the nucleoid, we provide evidence that the cytoplasm is a poor solvent for the chromosome in *Escherichia coli*. Monte Carlo simulations showed that such a poor solvent compacts the chromosome and promotes spontaneous formation of chromosomal domains connected by lower-density DNA regions. Cryo-electron tomography and fluorescence microscopy revealed that the (poly)ribosome density within the nucleoid is spatially heterogenous and correlates negatively with DNA density. These findings have broad implications to our understanding of chromosome folding and intracellular organization.

## Introduction

All cells, regardless of their origin, must package and organize their genome in a small volume. Proper folding of the chromosomes is critical as it affects many cellular processes, including gene expression, DNA repair and chromosome segregation. Unlike eukaryotic cells, bacterial cells lack a nuclear membrane and do not package their DNA content into repeating structural units akin to nucleosomes. However, they still fold and concentrate their chromosomal material into a dynamic and organized DNA meshwork known as the nucleoid. In many bacterial species, including *Escherichia coli*, the nucleoid does not spread throughout the cell; instead, it is found within a portion of the cytoplasmic space (Figure 1) (Gray et al., 2019; Hobot et al., 1985; Kellenberger et al., 1958; Mason and Powelson, 1956; Piekarski, 1937; Robinow and Kellenberger, 1994). This indicates that the compaction of the bacterial chromosome is not simply dictated by the physical confinement created by the cell envelope. Furthermore, the chromosome displays higher-order organization across multiple length scales (Dame et al., 2020; Verma et al., 2019). For instance, high-resolution chromosome conformation capture (Hi-C) studies in different bacterial species have demonstrated that the chromosome is organized into various “chromosomal interaction domains” (CIDs) within which nearby gene loci interact more frequently with each other than with those outside the domain (Le et al., 2013; Lioy et al., 2018; Marbouty et al., 2014; Marbouty et al., 2015; Val et al., 2016; Wang et al., 2017; Wang et al., 2015).

**Figure 1.**
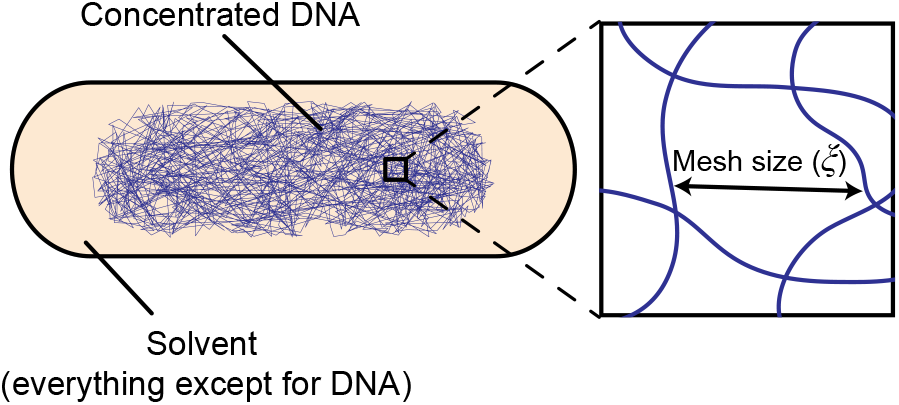
The *E. coli* cytoplasm viewed from a polymer physics perspective. Schematic showing the *E. coli* chromosome folded into a meshwork structure known as the nucleoid. The nucleoid mesh size is denoted as *ξ*. The cytoplasm can be viewed as a semidilute polymer solution, where the DNA is the polymer and the rest of the cytoplasm acts as its solvent.

Apart from its compacted and organized structure, another important, through less appreciated, aspect of the nucleoid is its mesh size (Figure 1), as it impacts the mobility of cytoplasmic components and thereby affects the spatial organization of the cytoplasm. For example, the enrichment of ribosomes/polyribosomes (mRNAs loaded with multiple ribosomes) at the cell poles and in-between segregated nucleoids in *E. coli* and other bacteria is often attributed to nucleoid exclusion (Azam et al., 2000; Bakshi et al., 2012; Gray et al., 2019; Lewis et al., 2000; Robinow and Kellenberger, 1994; Sanamrad et al., 2014). It is, however, difficult to predict which cytoplasmic components are impacted by the nucleoid, as its average mesh size has not been measured in any bacteria.

What are the biophysical principles that help explain the compacted size, domain organization and mesh size of the bacterial nucleoid? Multiple factors are thought to be at play (Surovtsev and Jacobs-Wagner, 2018). They include transcription and other biochemical processes that modulate DNA supercoiling (Dorman, 2019; Ma and Wang, 2016). Bacteria also carry a number of DNA-binding proteins, known as nucleoid-associated proteins (NAPs) and structural maintenance of chromosome (SMC) proteins, that alter the structure of the DNA locally or over long molecular distances (Dame et al., 2020; Dillon and Dorman, 2010). While the direct contribution of these proteins to DNA compaction is not entirely clear (Spurio et al., 1992; Wu et al., 2019), NAPs and SMC protein complexes are known to play important roles in the organization and regulation of chromosome architecture at the level of individual genes and CIDs (Dame et al., 2020). In addition, macromolecular crowding is frequently proposed to drive chromosome compaction through steric effects (Cunha et al., 2001; de Vries, 2010; Jeon et al., 2017; Jun, 2015; Odijk, 1998; Pelletier et al., 2012; Shendruk et al., 2015; Wegner et al., 2016; Wu et al., 2019; Yang et al., 2020; Yoshikawa et al., 2010; Zhang et al., 2009; Zimmerman, 1993; Zimmerman and Minton, 1993). The idea is that cytoplasmic components larger than the DNA mesh size (i.e., crowders) will be excluded from the nucleoid, creating an imbalance in component concentration between the nucleoid region and the rest of the cytoplasm. This imbalance results in an effective osmotic pressure that pushes DNA segments closer to each other. The magnitude of compaction driven by this steric repulsion is currently unknown inside cells.

It is important to note that the DNA and the cytoplasmic components do not interact only sterically, as neither of them are chemically inert. From a simple polymer physics perspective, the cytoplasm can be viewed as a polymer solution: the polymer is the DNA, and the solvent is everything else in the cytoplasm (water, metabolites, proteins, RNAs, etc.) (Figure 1). It is well established that the interaction between a polymer and its solvent directly affects the three-dimensional (3D) conformation of the polymer (Gennes, 1979; Rubinstein, 2003). Based on the polymer-solvent interaction, the quality of a solvent is broadly classified into three types: good, ideal and poor. In a good solvent, interactions between the polymer and the solvent are favored over interactions between polymer segments. Conversely, in a poor solvent, interactions between polymer segments are favored over their interactions with the solvent. When the repulsive and attractive interactions are balanced out (zero net interaction), the solvent is said to be ideal. Despite the well-known importance of the solvent for the polymer conformation, the effects of the solvent quality of the bacterial cytoplasm (or the eukaryotic nucleoplasm) on DNA compaction or organization are unclear. This is presumably because the solvent quality of the cytoplasm (or nucleoplasm) has not been measured and is difficult to predict. Traditional methods for determining the solvent quality of a polymer solution, such as dynamic light scattering, small-angle X-ray scattering, rheology, nuclear and magnetic resonance (Auge et al., 2009; Guettari et al., 2012; Waigh et al., 2001), are unfortunately not suitable for measurements inside cells. In this study, we develop an experimental approach to estimate the solvent quality of the cytoplasm in *E. coli* cells. Our results show that the cytoplasm is a poor solvent for the chromosome and that this macroscopic characteristic of the cytoplasm contributes to chromosome folding and intracellular organization.

## Results

As the polymer concentration increases, the polymer solution transitions from the regime of being dilute to semidilute, where the polymer segments overlap to form a meshwork. It is well established in polymer physics that the average mesh size (or correlation length) ξ of a semidilute polymer solution depends not only on properties of the polymer (concentration, rigidity, molecular weight and monomer size), but also on the quality of the solvent (Gennes, 1979; Rubinstein, 2003), as shown in Eq. 1 (see STAR Methods for derivation).

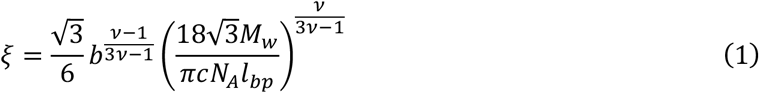

In this equation, the term *c* represents the average concentration of the polymer. The Flory exponent *v* quantifies the solvent quality. It is equal to 0.5 for an ideal solvent, and is smaller than 0.5 for a poor solvent and greater than 0.5 for a good solvent (Rubinstein, 2003). *N_A_* is Avogadro’s number. *M_w_* and *l_bp_* are the average molecular weight (650 g/mol) and size of a base pair (0.34 nm), respectively (Diekmann et al., 1982; Lee et al., 2012; Peale et al., 1989; Ratilainen et al., 2001; Yonemura and Maeda, 1982). The remaining term, *b*, is the Kuhn length of the polymer, which is a measure of its rigidity and equals to twice the persistence length of the polymer. DNA is a semiflexible polymer. Its rigidity stems from base stacking interactions within the DNA duplex and electrostatic repulsion among phosphate groups in the DNA backbone. The persistence length of the DNA is often assumed to be 50 nm (i.e., Kuhn length *b* = 100 nm). However, in vitro measurements of DNA rigidity indicate that the presence of multivalent cations, including Mg^2+^, the most abundant divalent cation in *E. coli* (Alatossava et al., 1985; Cayley et al., 1991; Kuhn and Kellenberger, 1985; Lusk et al., 1968; Moncany and Kellenberger, 1981), can decrease the DNA persistence length down to 25-35 nm (Baumann et al., 1997; Mantelli et al., 2011; Porschke, 1986; Porschke, 1991). Based on these measurements and the millimolar concentration of Mg^2+^ in the *E. coli* cytoplasm (Alatossava et al., 1985; Cayley et al., 1991; Lusk et al., 1968), the Kuhn length of the chromosomal DNA (Ď) is estimated to be ~60 nm (i.e., a persistence length of ~30 nm).

Given the aforementioned values for *b*, *M_w_* and *l_bp_*, the solvent quality of the cytoplasm can be deduced using Eq. 1 by determining the concentration of the chromosomal DNA within the nucleoid region (*c*) and the average mesh size of the nucleoid (*ξ*).

### Average DNA concentration within the nucleoid

To determine the average concentration of DNA within the nucleoid, we took advantage that in nutrient-poor environments (e.g., glycerol as a carbon source), *E. coli* cells exhibit discrete B, C and D periods, corresponding to the cell cycle phases before, during and after DNA replication, respectively (Cooper and Helmstetter, 1968). In B-period cells, the DNA mass can be unambiguously determined, as each nucleoid in these cells consists of a single, non-replicating chromosome (Cooper and Helmstetter, 1968). Using a previously described method (Gray et al., 2019), we identified B-period cells within a population by examining cell size and the spatial pattern of the DNA replication marker SeqA fused to mCherry. During DNA replication (C period), SeqA binds the newly synthesized hemimethylated DNA (Brendler et al., 1995; Lu et al., 1994; Slater et al., 1995; Waldminghaus et al., 2012). This property leads to the formation of fluorescent foci in cells actively replicating DNA (Figure 2A), as shown before (Adiciptaningrum et al., 2015; Gray et al., 2019; Helgesen et al., 2015; Molina and Skarstad, 2004; Wallden et al., 2016). In contrast, SeqA-mCherry displayed a diffuse distribution in cells before and after DNA replication (i.e., during B and D periods, respectively) (Figure 2A). We used these changes in spatial distribution of SeqA-mCherry signal and the knowledge that cells grow in size during the cell cycle to classify cells by cell-cycle period (Figure 2B). Next, we identified the contour of DAPI-stained nucleoids in 19,510 cells in the B period using the objectDetection module of the Oufti software package (Paintdakhi et al., 2016) and quantified their volumes (STAR Methods). We found that nucleoids consisting of one chromosome have an average volume of ~0.7 μm^3^ (Figure 2C). The mass of the *E. coli* chromosome is about 5×10^-12^ mg given that this chromosome is made of ~4.6×10^6^ base pairs and that the average molecular weight of a base pair is 650 g/mol (Lee et al., 2012; Peale et al., 1989; Ratilainen et al., 2001). From this, we determined the average DNA concentration within the nucleoid region to be 7.1 ± 1.0 mg/ml (Figure 2D). Our measurements, which were done with cells growing at 37°C, were robust to a change in growth temperature to 30°C (Figure S1).

**Figure 2.**
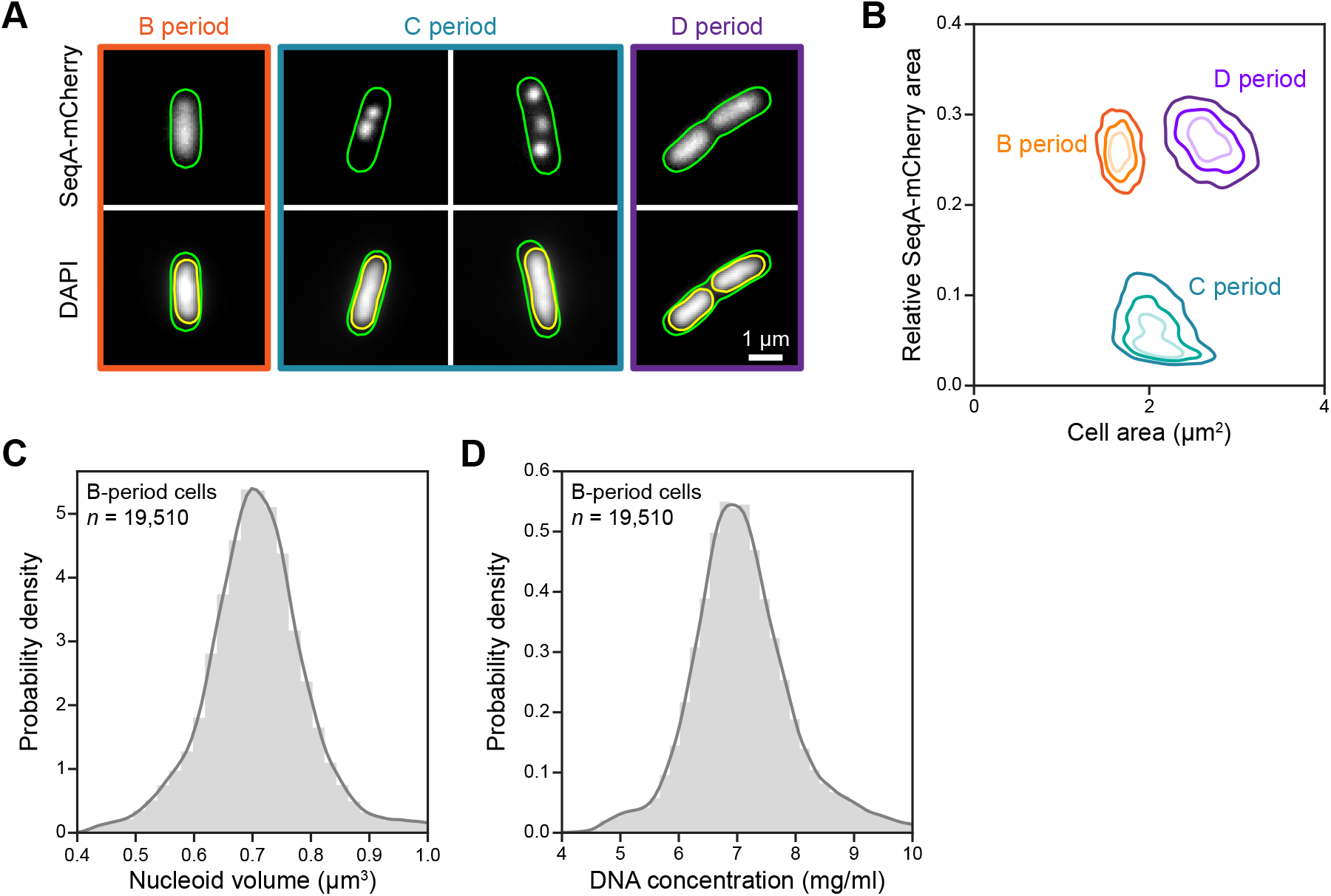
Estimation of the average DNA concentration in the nucleoid region in B-period *E. coli* cells. A. Fluorescence images of representative DAPI-stained CJW6324 cells in different cell cycle periods: B, C and D periods (i.e., before, during and after the DNA replication). The green and yellow outlines are the cell and nucleoid contours detected by the software package Oufti. B. Plot showing the relative area of the SeqA-mCherry signal as a function of cell area. The contour lines (from light to dark color) represent the 25%, 50% and 75% probability envelopes of the data. C. Probability density function of the estimated nucleoid volume in B-period cells. D. Probability density function of the average DNA concentration calculated based on the total mass of the *E. coli* chromosome and the nucleoid volume measurements shown in panel C. See also Figure S1.

Under nutrient-rich (fast growth) conditions, *E. coli* cells enlarge to accommodate multi-fork DNA replication. The absence of a discrete B period under these conditions, prevented us from performing the same microscopy analyses as above. However, when we converted bulk measurements of DNA mass per cell made under nutrient-rich growth conditions (Basan et al., 2015) to DNA concentrations within the nucleoid volume (*c*) by taking into consideration of changes in cell and nucleoid volume, we obtained *c* = 7.4 ± 0.2 mg/ml (*n* = 3 nutrient-rich growth conditions, see STAR Methods for details). The agreement between this estimation and our single-cell measurements (Figure 2D) indicates that *c* remains largely constant across nutrient-poor and -rich conditions. This is consistent with both DNA content per cell and nucleoid size scaling with cell size across growth conditions (Basan et al., 2015; Gray et al., 2019; Sharpe et al., 1998).

### Apparent average nucleoid mesh size in *E. coli* cells

Given *c* ≈ 7 mg/ml, apart from the Flory exponent (*v*), the remaining unknown in Eq. 1 is the mesh size (*ξ*). Studies on biopolymer solutions and hydrogels have demonstrated that particles of size much smaller than the average mesh size diffuse freely through the meshwork (Axpe et al., 2019; Cai et al., 2011; Carn et al., 2012; Wong et al., 2004). When the probe size is close to the average mesh size, the dynamics of the probes becomes impacted by the polymer mesh. This impact exacerbates as the probes increase in size. In the context of the bacterial cell, this implies that probes much smaller than the nucleoid mesh size will diffuse freely in the cytoplasm without any apparent hindrance from the nucleoid (assuming that the particles do not interact with the nucleoid). Such probes will have an equal probability of diffusing inside or outside the nucleoid. In contrast, probes larger than the average mesh size will have a higher probability of diffusing outside the nucleoid, i.e., nucleoid exclusion will increase with increasing probe size.

If we assume that the cytoplasm is an ideal solvent (*v* = 0.5) for the chromosome, the average mesh size should be 22 nm (Eq. 1), in which case particles larger than 22 nm should have their motion and spatial distribution affected by the nucleoid. To test this assumption experimentally, we used an *E. coli* strain (CJW6340) that expresses a GFP-tagged, artificially designed protein that self-assembles into a 60-subunit dodecahedron nanocage of 25 nm in diameter (Figure 3A) (Hsia et al., 2016). These nanocages were originally designed for drug and vaccine delivery applications. However, because of their synthetic nature, they are foreign to the bacterial cytoplasm and therefore unlikely to form specific interactions with the nucleoid or other cytoplasmic components, allowing us to repurpose them as intracellular tracers.

**Figure 3.**
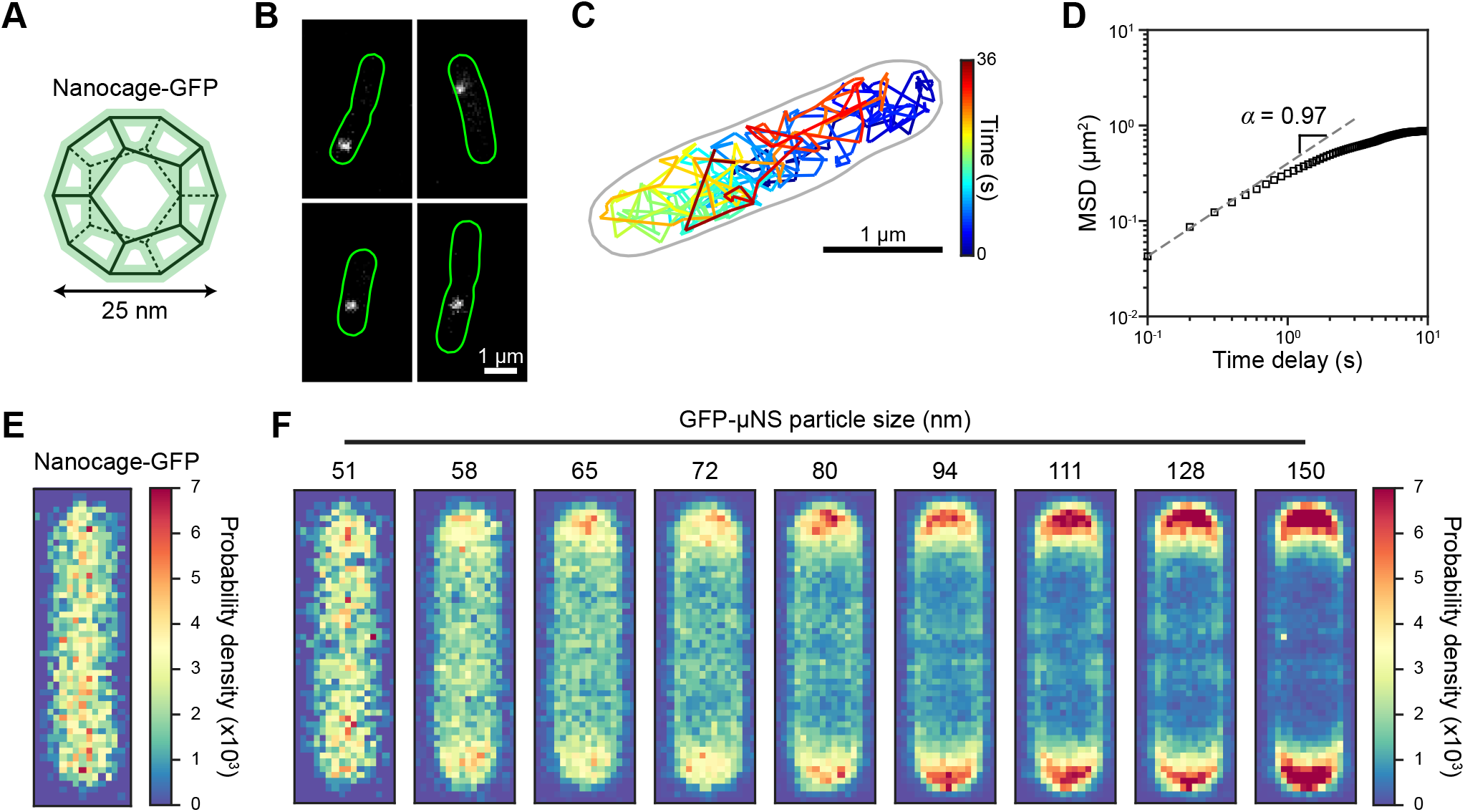
Estimation of the average nucleoid mesh size. A. Schematic of an assembled nanocage-GFP particle. The particle has a dodecahedron shape (Hsia et al., 2016). B. Fluorescence images of nanocage-GFP particles assembled in *E. coli* cells (CJW6340). C. Example of nanocage-GFP particle trajectory. The gray outline indicates the cell contour detected by the Oufti software package using the corresponding phase image of the cell. D. Log-log ensemble-averaged mean squared displacement (MSD) based on individual trajectories of nanocage-GFP particles (*n* = 85). The slope (*α* = 0.97) was fitted based on the first three time delays. E. Probability density map of the localization of nanocage-GFP particles. The density map was constructed by plotting the normalized 2D histogram of the relative particle positions. F. Probability density maps of the localization of GFP-μNS particles as a function of their size. The positions of the particles (*n* = 133,692) were determined relative to the cell contours. Particles were then binned by their sizes, as indicated. For each particle-size bin, a probability density map of particle localizations was constructed, as described in panel E. See also Figure S2 and Video S1.

Basal expression of nanocage-GFP proteins (i.e., leaky synthesis) was sufficient to produce a single nanocage-GFP particle per cell in a fraction of the population (Figure 3B). Single-particle tracking experiments revealed no evidence of steric hindrance by the nucleoid, as the nanocage-GFP particle appeared to diffuse freely throughout the cytoplasm (Figure 3C; Video S1). This was further supported by calculating the ensemble-averaged mean squared displacement (MSD) based on all particle trajectories measured (*n* = 85). At the short time scale (first three points), the MSD varied linearly with the time delay, as the slope of MSD curve in the log-log scale was close to 1 (Figure 3D). This indicates that the diffusion of nanocage-GFP particles was Brownian (at the long time scale, the MSD deviated from the linearity due to cell confinement). Furthermore, ensemble calculation of the relative positions of nanocage-GFP particles inside cells demonstrated that the probability density of nanocage-GFP localization is uniform throughout the cytoplasm (*n* = 2,500 localizations) (Figure 3E). Thus, these 25-nm probes were neither trapped nor excluded by the nucleoid, consistent with their diffusion being Brownian. This result indicates that the nucleoid mesh size is greater than 25 nm, which is inconsistent with the bacterial cytoplasm being an ideal solvent for the chromosome.

To estimate the average nucleoid mesh size, we used GFP-μNS particles, which are larger probes that our laboratory previously developed to probe the material properties of the cytoplasm (Parry et al., 2014). GFP-μNS is a fluorescent protein fusion to a mammalian reovirus protein that self-assembles into a globular complex (Broering et al., 2005; Broering et al., 2002). Like GFP and the nanocage protein, μNS is not of bacterial origin and is therefore unlikely to display a significant affinity to components of the *E. coli* cytoplasm. When expressed from the chromosome of *E. coli* (CJW4617) following IPTG induction (50-200 μM) for 30 to 120 min (STAR Methods), GFP-μNS assembled into fluorescent particles, usually one per cell (Parry et al., 2014). The size of the GFP-μNS particles, which was deduced by establishing a calibration between the particle sizes and their corresponding fluorescence intensities (Parry et al., 2014) (STAR Methods), ranged from ~50 to ~200 nm based on the level of protein expression in each cell (Figure S2).

We tracked GFP-μNS particles (*n* = 133,692) and calculated their relative positions in the corresponding cells. We then binned all GFP-μNS particles by size and, for each size bin, constructed a probability density map of the relative particle position in the cells (Figure 3F). These probability density maps indicate the likelihood of finding a GFP-μNS particle at a specific intracellular location across cells. For particles in the bin with the smallest average particle size (51 nm), the probability density map appeared indistinguishable from that obtained for nanocage-GFP particles (Figure 3E), indicating that these particles are not excluded by the nucleoid. This pattern changed in the next size bin, as probes of an average size of 58 nm started to display an increased probability to be localized at the cell pole regions (i.e., away from the nucleoid). This enrichment in probability density of localization at the cell poles continued to increase with probe size (Figure 3F), with ever larger particles becoming less and less likely to be found within the nucleoid region. Particles with the largest average size (150 nm) were almost completely excluded from the nucleoid, as shown by the high probability density of localization at the cell poles and the near-zero probability density elsewhere in the cell. While these ensemble results cannot provide an exact value, they approximate the apparent average nucleoid mesh size to be around 50 nm.

### The *E. coli* cytoplasm behaves as a poor solvent for the chromosome

Based on *c* ≈ 7 mg/ml, *b* ≈ 60 nm and *ξ* ≈ 50 nm, we found that the corresponding Flory exponent, *v*, is ~0.36 (Eq. 1). Such a small Flory exponent (< 0.5) implies that the bacterial cytoplasm is a poor solvent for the chromosome. Our conclusion is robust against variability in DNA concentration across cells (Figure 2D) or nutrient-poor or -rich growth conditions (see above). For instance, a DNA concentration of 8 mg/ml would correspond to an even smaller Flory exponent (Figure 4A). At DNA concentrations of 6 mg/ml, or even as low as 5 mg/ml, the Flory exponent, *v*, remains well below 0.5, consistent with the cytoplasm behaving as a poor solvent.

**Figure 4.**
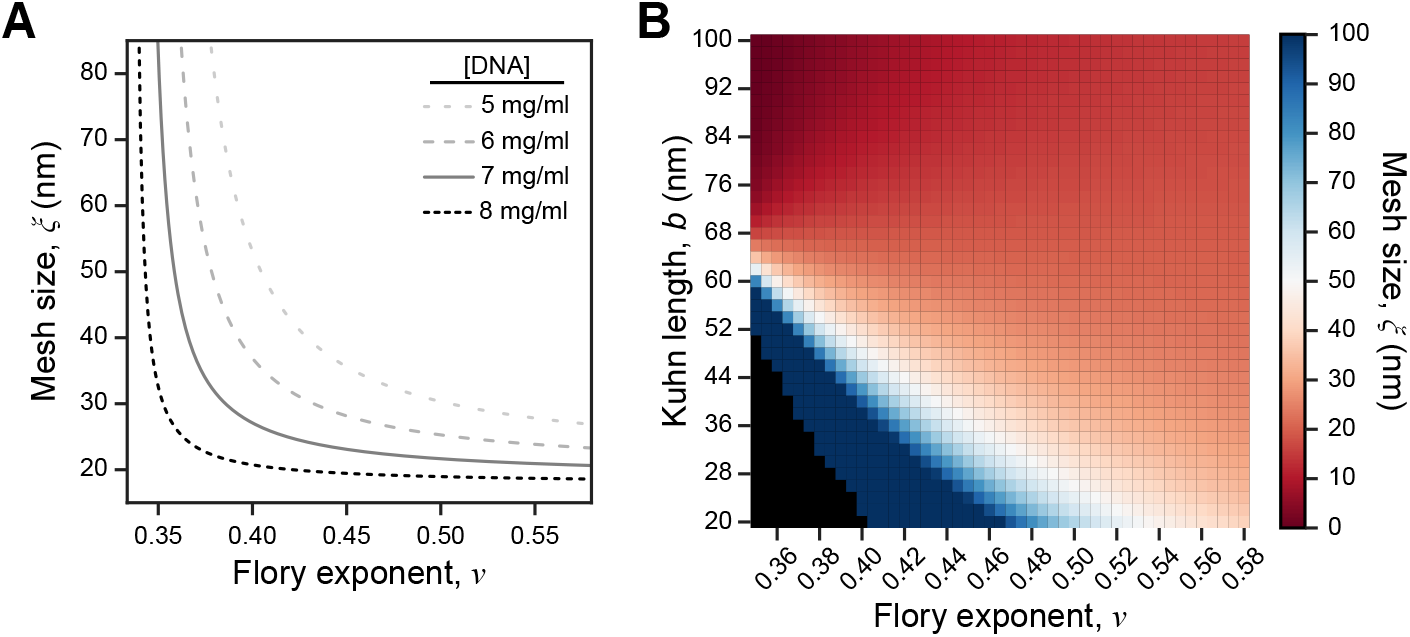
Relationship between solvent quality, polymer concentration, Kuhn length and mesh size. A. Plot showing the average mesh size of the nucleoid as a function of the Flory exponent, with a DNA Kuhn length of 60 nm and various DNA concentrations (5, 6, 7 or 8 mg/ml) calculated using Eq. 1. B. Heatmap showing the average nucleoid mesh size calculated using Eq. 1 with different combinations of DNA Kuhn lengths and Flory exponents, at a given DNA concentration of 7 mg/ml.

The phase diagram in Figure 4B shows that our conclusion is also robust to variations in DNA Kuhn length due to potential fluctuations in cytoplasmic ionic strength. To achieve the observed apparent ~50-nm nucleoid mesh size in an ideal solvent (*v* = 0.5) or a good solvent (e.g., *v* = 0.58), the Kuhn length of DNA at an average concentration of 7 mg/ml would have to be 26 nm or 13 nm, respectively (Eq. 1). This would mean a DNA persistence length (half the Kuhn length) equal to 13 nm or lower. Such a small value has never been observed experimentally even in solutions of extremely high ionic strength (e.g., 3-4 M NaCl) (Borochov et al., 1981; Kam et al., 1981; Sobel and Harpst, 1991) or in the presence of multivalent ions such as Mg^2+^ (Baumann et al., 1997; Mantelli et al., 2011; Porschke, 1986; Porschke, 1991).

### A poor solvent promotes the organization of the chromosome into domains

Intriguingly, for a DNA Kuhn length of 60 nm and an average DNA concentration of 7 mg/ml, Eq. 1 indicates that the average mesh size decreases as the Flory exponent increases (i.e., as the solvent quality improves) (Figure 4A). To understand how the solvent quality of the cytoplasm affects the nucleoid mesh size at the given DNA concentration and Kuhn length, we performed 3D Monte Carlo simulations of chromosome conformation in poor (*v* = 0.36), ideal (*v* = 0.50) and good (*v* = 0.58) solvents (STAR Methods). For all simulations, we modeled the entire *E. coli* chromosome (4.6 million base pairs, contour length ≈ 1.6 mm) as a chain of 26,066 segments, with each segment having a length of 60 nm (corresponding to the Kuhn length). To achieve the average DNA concentration observed in actual nucleoids (7 mg/ml), we confined the entire chromosome within a spherocylindrical space with a volume (0.70 μm^3^) equal to the experimentally determined average nucleoid volume in B-period cells (Figure 2C).

The simulations revealed a drastic difference in chromosomal conformation when varying the solvent quality (Video S2). The DNA density in the poor solvent appeared to be much more spatially heterogeneous than that in the ideal or good solvent. This was also evident in cross-sectional slices of simulated chromosomes in which each dot represents a DNA segment crossing the plane (Figure 5A). The spatial heterogeneity of DNA density in the poor solvent condition compared to the other conditions was also apparent in the two-dimensional (2D) histograms showing the probability density of finding a DNA segment inside subregions of the nucleoid (Figure 5B). This result is consistent with the spatial heterogeneity of DNA density observed in super-resolution fluorescence images of bacterial nucleoids (Le Gall et al., 2016; Marbouty et al., 2015; Spahn et al., 2014; Spahn et al., 2018; Stracy et al., 2015).

**Figure 5.**
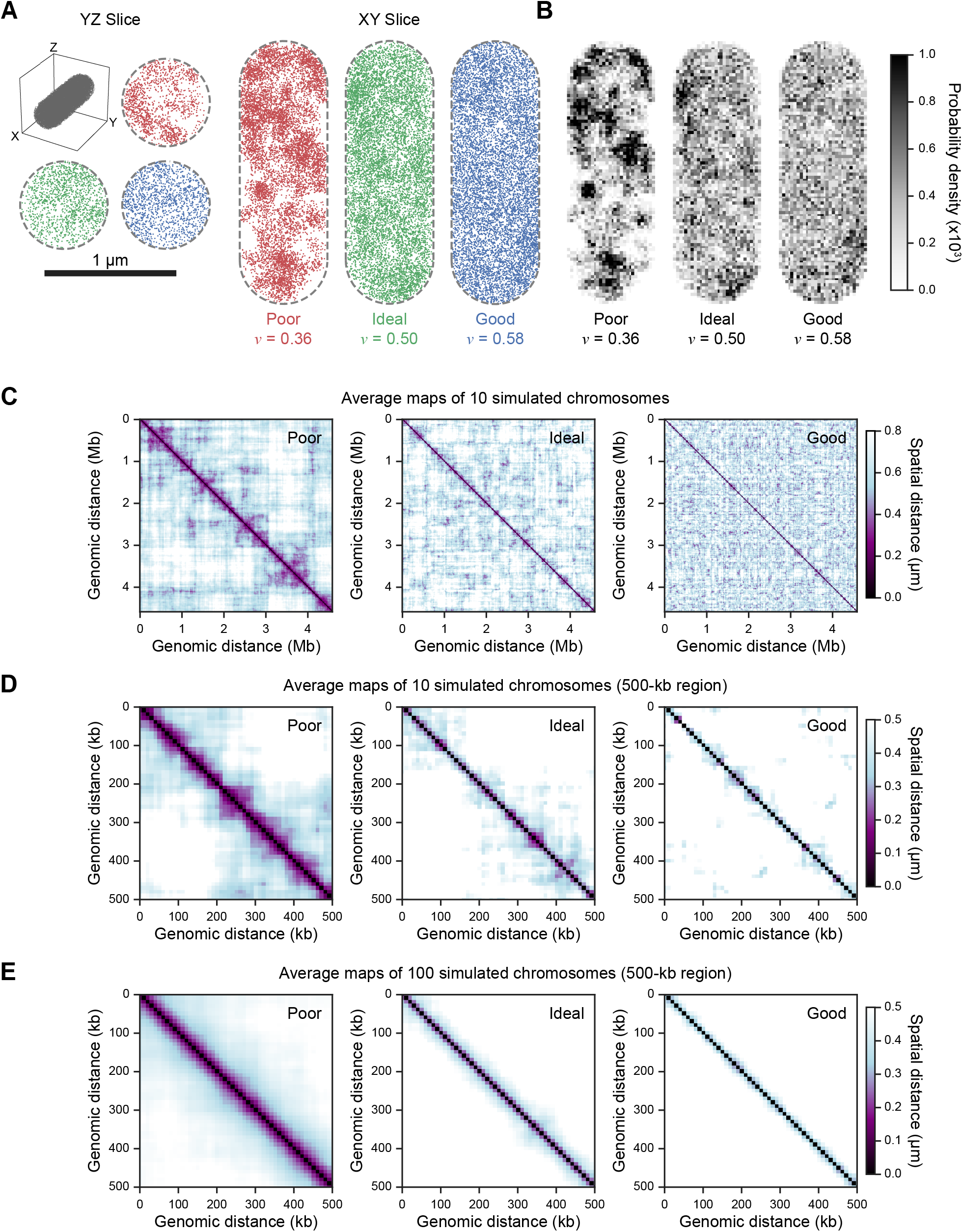
Monte Carlo simulations of the *E. coli* chromosome conformation in different types of solvent. The entire *E. coli* chromosome was modeled using Monte Carlo simulations (STAR Methods). Chromosomes were simulated within the confinement of a spherocylinder with a volume equal to that of an average nucleoid (~0.7 μm^3^) in a poor (*v* = 0.36), ideal (*v* = 0.50) and good solvent (*v* = 0.58). The Kuhn length of the DNA was assumed to be 60 nm. A. Example slices of simulated chromosomes in a poor, ideal or good solvent. Each dot represents a DNA segment going in or out of the slices (XY- or YZ-plane). B. Density maps of the slices shown in panel A. The maps were constructed as the normalized 2D histograms of positions of the simulated DNA segments. C. Distance maps indicating the spatial separation between any two loci along the simulated chromosomes in the poor, ideal or good solvent. Each map is a result of averaging 10 simulated chromosomes. D. Distance maps as in panel C, except for showing only the first 500-kb region of the simulated chromosomes. E. Distance maps as in panel D, except that 100 simulated chromosomes were averaged instead of 10. See also Figure S3 and Video S2.

In the poor solvent simulations, DNA segments were locally attracted to each other, leading to the formation of regions with high DNA density interspersed with regions of low DNA density. This spatial heterogeneity in DNA density created “holes” (Figure 5A; Video S2), which may increase the apparent average mesh size and allow larger objects to pass through the nucleoid. Meanwhile, the DNA density of the nucleoid was more spatially homogenous in the ideal and good solvents (Figure 5A; Video S2), likely contributing to the smaller apparent average mesh size.

The denser, domain-like regions of the nucleoid in the poor solvent (Figure 5A-B; Video S2) reminded us of CIDs reported in Hi-C studies in *E. coli* and other bacteria (Le et al., 2013; Lioy et al., 2018; Marbouty et al., 2014; Marbouty et al., 2015; Val et al., 2016; Wang et al., 2017; Wang et al., 2015). To examine this more closely, we calculated the Euclidean distance between pairs of DNA loci along simulated chromosomes binned by 10-kilobase pairs. From this information, we created average maps that indicate the distances between the pairs of DNA loci along the entire simulated chromosomes or a 500-kb region under a poor, ideal or good solvent condition (Figure 5C-D). Along the diagonal of the distance maps, the distance is always 0 (black), because the separation between any given DNA locus and itself is zero by definition. Dark patches along the diagonal in the distance maps represent individual domain-like structures, in which DNA loci that extend a relatively large genomic distance remain in close spatial proximity to each other. Large domain-like structures were only seen under the poor solvent condition (Figure 5C-D). Thus, by itself, the poor solvent quality of the cytoplasm results in the formation of chromosomal domains that are reminiscent of CIDs seen in Hi-C experiments.

However, the poor solvent quality of the cytoplasm alone cannot explain *where* these domains form. In our simulations, the DNA has no sequence information. Therefore, the formation of domains can occur anywhere along the DNA polymer, as shown by randomly averaging 10 different simulated chromosomes (Figure S3A). In Hi-C experiments, results are based on the average of billions of cells. In our case, averaging more simulated chromosomes (e.g., 100) resulted in the disappearance of distinct domain boundaries in distance maps (Figure 5E and Figure S3B). Interestingly, Hi-C experiments on rifampicin-treated *Caulobacter crescentus* cells show that global inhibition of transcription results in the dissolution of most domain boundaries (Le et al., 2013). Collectively, our results suggest that the poor solvent quality of the cytoplasm promotes domain formation, but DNA sequence-dependent factors (e.g., transcription) are responsible for setting the domain boundaries at consistent chromosomal locations across cells (see Discussion).

### Spatial heterogeneity in ribosome density within the nucleoid correlates negatively with DNA density

With the poor solvent quality of the cytoplasm resulting in spatial heterogeneity of DNA density, we reasoned that this may, in turn, affect the spatial distribution of other cytoplasmic components, even within the nucleoid region. Specifically, we hypothesized that ribosomes (which exist mostly in polyribosome form) are not only enriched outside of the nucleoid region (Azam et al., 2000; Bakshi et al., 2012; Gray et al., 2019; Robinow and Kellenberger, 1994; Sanamrad et al., 2014), but also heterogeneously distributed throughout the nucleoid due to its uneven DNA density. To test this hypothesis, we prepared frozen-hydrated cryo-electron tomography (cryo-ET) samples of exponentially growing *E. coli* cells (MG1655) (Figure 6A, STAR Methods). To visualize the native *E. coli* cytoplasm with higher contrast and resolution, we used cryo-focused ion beam (cryo-FIB) milling to produce lamellae of a thickness between 150 and 260 nm (Figure 6A and Figure S4A-B). Lamella reconstruction revealed ribosomes as dark spots distributed within the *E. coli* cytoplasm (Figure 6B; Video S3).

**Figure 6.**
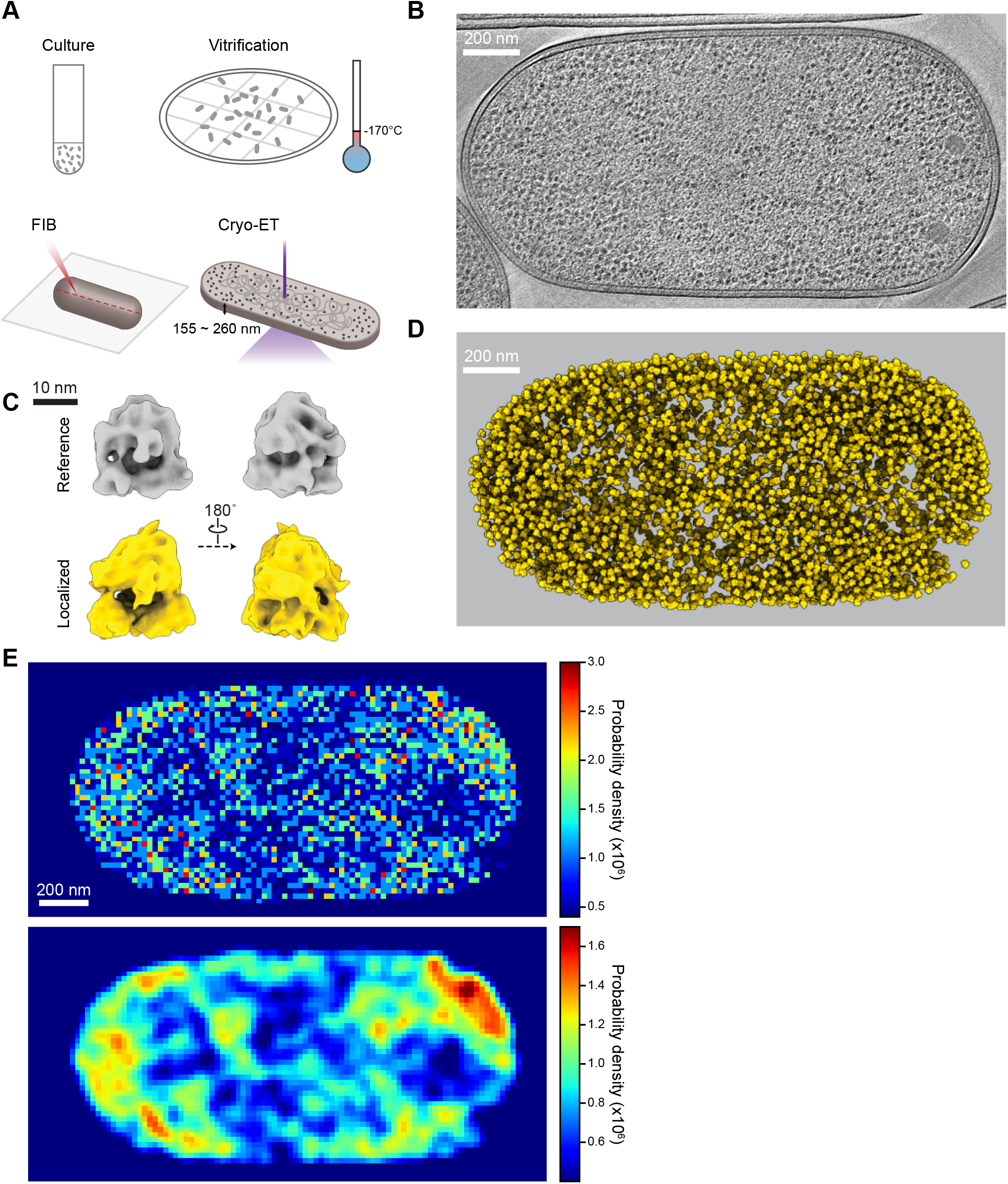
Spatial distribution of ribosomes in *E. coli* tomograms. A. Schematic showing the preparation procedure for the cryo-ET samples. Wild-type *E. coli* cells (MG1655) were harvested in exponential phase, concentrated by centrifugation, deposited on the grid and vitrified in liquid ethane. Using cryo-FIB milling, a thin lamella sample of thickness between 155 and 260 nm was obtained for the cryo-ET imaging. B. Example section of the tomogram. C. Comparison between the reference ribosome structure (gray) and the averaged structure (yellow) derived from ribosomes localized in the lamella tomogram shown in panel B. An *E. coli* 70S ribosome structure (EMD-20173) was used as the reference (Fu et al., 2019). See also Video S4 for a 3D overview. D. Maps showing localized ribosome structures (yellow) with the positions and orientations determined in the tomogram shown in panel B. E. Probability density maps of ribosome structures localized in the tomogram in panel B. The top represents the probability density map of the localizations of ribosomes determined from the tomogram. The map was constructed as a 2D histogram of the ribosome positions in the XY-plane. The bottom map was obtained after Gaussian smoothing (*σ* = 1.2 pixel, 21 nm) of the top map to more clearly show subcellular regions with high and low density in ribosomes. See also Figure S4, Video S3 and Video S4.

To precisely localize the ribosomes, we used the template search routines in emClarity (Himes and Zhang, 2018) to obtain 3D positions and orientations of the ribosomes detected in the lamella tomograms. Sub-volumes of the detected ribosomes were aligned and classified (STAR Methods, Figure S4C-D) (Winkler, 2007; Winkler et al., 2009). The average ribosome structure calculated using the detected ribosomes was similar to that of the low-pass filtered reference structure (Figure 6C; Video S4) (Fu et al., 2019), suggesting a high degree of accuracy in our detection procedure. In total, we detected 5,028 ribosomes in the tomogram represented in Figure 6D. Based on the volume of the lamella (0.21 μm^3^), the average ribosome density in the tomogram was approximately 24,000 μm^-3^, which is consistent with the reported ribosome density under similar growth conditions (Bakshi et al., 2012; Bremer and Dennis, 2008). We found that the density of (poly)ribosomes was highly heterogeneous across the lamella. This was readily apparent after the Gaussian smoothing of the ribosome density map (Figure 6E and Figure S4E). In addition to the expected accumulation of (poly)ribosomes at the cell periphery (i.e., away from the nucleoid region), (poly)ribosomes displayed heterogeneity in density across the cytoplasm, including within the expected nucleoid region (i.e., central region of the cell).

Since DNA cannot be directly visualized in the tomograms, we turned to fluorescence microscopy to examine whether the spatial heterogeneity in ribosome density within the nucleoid region is linked to differences in DNA densities. We imaged DAPI-stained *E. coli* cells (CJW7020) producing 50S ribosomal protein L1 tagged with monomeric superfolder GFP (Figure 7A-B). As before, in each cell (*n* = 1,126), we identified the nucleoid outlines based on the DAPI signal (dotted lines in Figures 7A-B). When calculating the correlation between the DNA and ribosome fluorescence signals, we considered only image pixels well within the nucleoid region—at least two pixels away from the nucleoid outline (third row images in Figure 7A-B)— to prevent any bias in the correlation analysis due to the well-known enrichment of (poly)ribosomes outside the nucleoid region. We then calculated the correlation coefficient (Spearman’s *ρ*) between the fluorescence signals of DAPI (DNA) and L1-msfGFP (ribosome) at the single-pixel resolution. We found that even well within the nucleoid, the ribosome fluorescence signal correlates negatively with the DNA signal, as shown with two cell examples (Figure 7C-D) as well as at the population level (*n* = 1,126 cells, Figure 7E). These findings are consistent with the idea that the solvent quality of the cytoplasm contributes to the uneven DNA density, which, in turn, affects (poly)ribosome localization (or vice versa, see Discussion).

**Figure 7.**
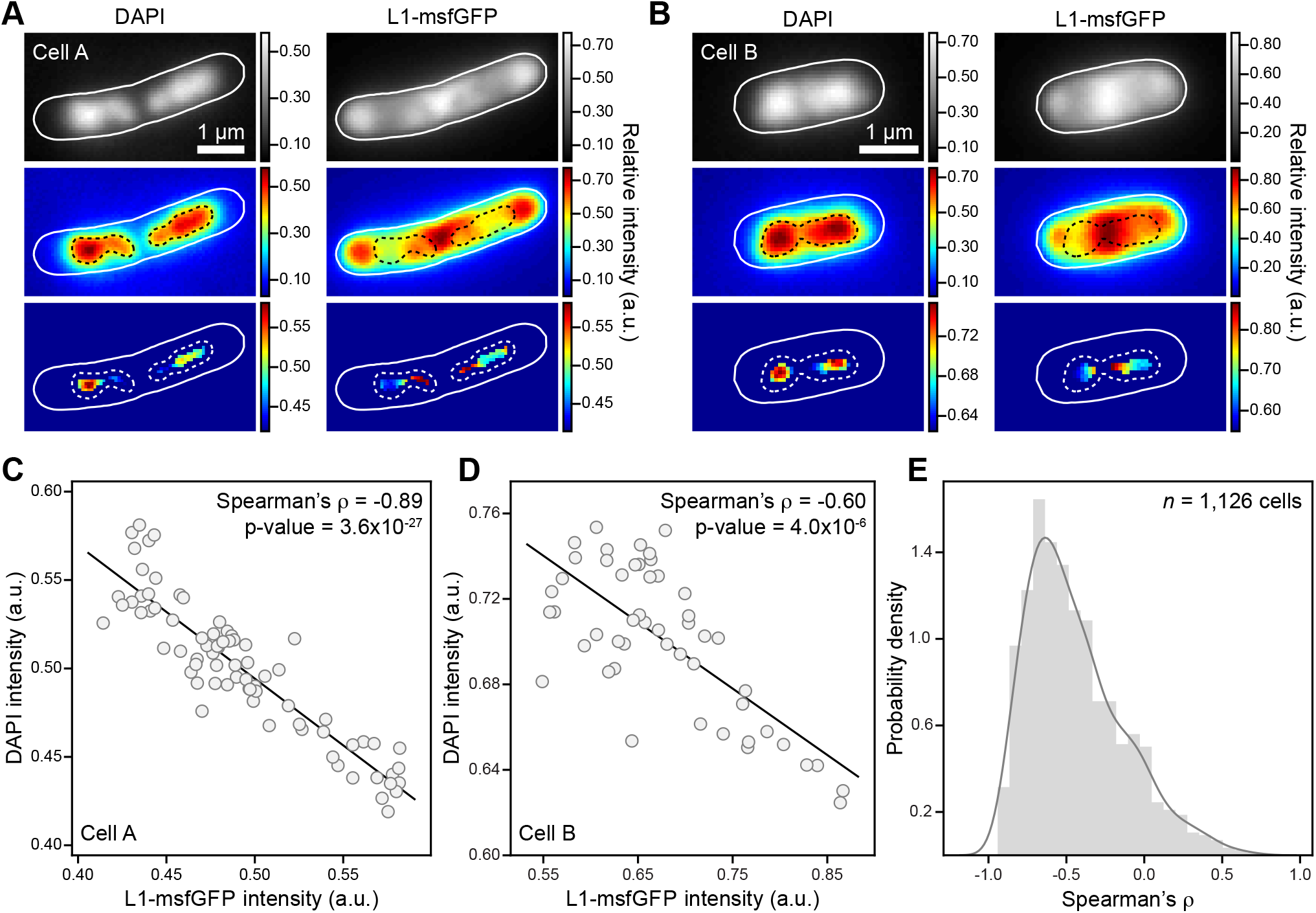
Pixel intensity correlation between DNA and ribosome fluorescence signals. A. Fluorescence micrographs showing an example cell (CJW7020) with a strong negative correlation (Spearman’s *ρ* = −0.89) between the DNA and ribosome fluorescence signals. The cell outlines are depicted as white solid lines, whereas the nucleoid outlines are shown as either black or white dash lines. The images are shown using either the conventional gray scale or a color scale. The bottom row shows only the pixels enclosed well within the nucleoid, at least two pixels away from the nucleoid outline (STAR Methods). These pixels were used for the correlation calculation. B. Similar to panel A, except for showing an example cell with a more common negative correlation (Spearman’s *ρ* = −0.60) between DNA and ribosome fluorescence signals. C. Scatter plot showing the pixel intensity correlation between DAPI and L1-msfGFP signals obtained in the cell shown in panel A. D. Similar to C, except for showing the corresponding pixel intensity correlation in the cell shown in panel B. E. Probability density function of pixel intensity correlations calculated among the cell population (*n* = 1,126).

### The poor solvent quality of the cytoplasm contributes to significant chromosome compaction

In addition to the internal mesh size and domain organization, compaction is another biophysical characteristic associated with the bacterial chromosome. A polymer is intrinsically more compact in a poor solvent because polymer segments preferentially interact with themselves rather than with the solvent. This fact prompted us to examine the level of compaction that the poor solvent quality of the cytoplasm could impose on the nucleoid. In our previous Monte Carlo simulations, the chromosome was modeled within the experimentally determined average volume of the nucleoid region (0.7 μm^3^) to keep the DNA concentration constant (i.e., 7 mg/ml) across solvent types. We performed another set of Monte Carlo simulations, but this time in an unbounded space to examine how the quality of the solvent affects the compaction of the chromosome (STAR Methods). The simulation results showed that the chromosome is most compact in the poor solvent condition, while being the most expanded in the good solvent (Video S5). The ideal solvent was associated with an intermediate phenotype.

To quantify the size of each simulated chromosome, we calculated its radius of gyration (STAR Methods). In these calculations, we took into consideration the circular geometry of the bacterial chromosome. Given a fixed contour length, the geometric constraint of being circular forces the chromosome to adopt a smaller average size compared to that of a linear chain. Quantitatively, the average radius of gyration of a circular polymer is a factor of 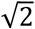 smaller than that of its linear counterpart (Casassa, 1965; Kramers, 1946; Rubinstein, 2003; Zimm and Stockmayer, 1949). We found that the volume of a circular DNA polymer of the same contour length as the *E. coli* chromosome is, on average, ~9 μm^3^ in a poor solvent (*v* = 0.36) compared to ~100 μm^3^ in an ideal solvent (*v* = 0.50) or ~550 μm^3^ in a good solvent (*v* = 0.58) (Figure 8). Thus, the poor solvent quality of the cytoplasm results in an additional ~10- and ~60-fold chromosome compaction, compared to the ideal and good solvent, respectively. This result suggests that the poor solvent quality of the cytoplasm plays a large role in chromosome compaction.

**Figure 8.**
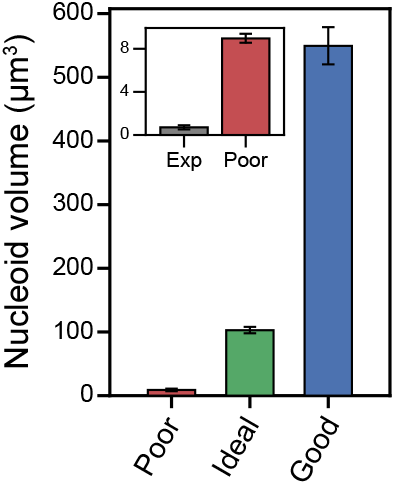
Chromosome compaction in different types of solvent. Bar graph showing the mean volume of 1,000 simulated chromosomes in each type of solvent. The error bars represent the 95% confidence interval of the mean. The inset shows a comparison between the volume of the nucleoid observed in the B-period *E. coli* cells (see Figure 2C) and the volume calculated for simulated chromosomes (*n* = 1,000) in the poor solvent. See also Video S5.

We note that this solvent effect alone cannot explain the full compaction of a single chromosome inside cells, as the actual volume of the nucleoid in B-period *E. coli* cells (Figure 2C) is still around 10 times smaller (Figure 8, inset). This is consistent with other compacting factors (e.g., DNA supercoiling, NAPs and macromolecular crowding) playing important roles as well (see Discussion).

## Discussion

The conformation of a given polymer depends on the quality of its solvent (Gennes, 1979; Rubinstein, 2003), yet, the solvent quality of the cellular milieu for the chromosome has been long overlooked, most likely due to the lack of a suitable method for its measurement inside cells. In this study, we developed such a method by taking a simple polymer physics perspective of the bacterial cytoplasm as a semidilute polymer solution, in which the chromosome is the polymer and its solvent is comprised of everything else in the cytoplasm. Using this approach, we found that the *E. coli* cytoplasm is a poor solvent for the chromosome and that this physicochemical property inherently interconnects three different aspects of the chromosome inside cells: compaction, mesh size and domain organization.

### Chromosome compaction

A significant level of compaction has long been considered mandatory for the bacterial chromosome to fit inside the cell. This was first suggested by early electron microscopy studies, where the chromosomes released from lysed cells were found to enlarge many times over the size of the cell (Kavenoff and Bowen, 1976; Kavenoff and Ryder, 1976). Dilution of NAPs and loss of macromolecular crowding are likely to contribute to this large nucleoid expansion, as often proposed. However, the solvent for the chromosome also drastically changed in these experiments, from the cytosol—a complex mixture of cytoplasmic components—to a simpler aqueous solution mostly composed of buffering reagents. Any considerable change in the chemical nature of a solvent is expected to affect the solvation and conformation of a dissolved polymer. In light of our finding that the cytoplasm behaves as a poor solvent for the chromosome, it is likely that the swelling of the chromosome following cell lysis is, at least in part, due to the release of the chromosome into a better solvent. Our work shows that in the cytoplasm, the chromosome is more likely to interact with itself compared to the rest of the cytoplasm. In such a poor solvent, preferential interaction (i.e., attraction) between parts of the chromosome compacts the entire chromosome ~10 to ~60 times more than if it were dissolved in an ideal or good solvent, respectively (Figure 8; Video S5).

Our finding also provided an opportunity for us to more carefully estimate how much a polymer of the contour length of the *E. coli* chromosome would need to be compacted to fit into the nucleoid region of an *E. coli* cell. Previous back-of-the-envelope calculations by different groups (Bloomfield, 1997; Holmes and Cozzarelli, 2000; Joyeux, 2014; Jun, 2015; Trun and Marko, 1998; Verma et al., 2019), including ours (Surovtsev and Jacobs-Wagner, 2018), suggested a compaction on the order of 1000-fold. These estimations relied on three assumptions. (i) Without prior knowledge, the bacterial cytoplasm was assumed to be an ideal solvent for the chromosome. As such, the chromosome was treated as an *ideal* chain that assumes the conformation of a random walk. In other words, the net interaction between the chromosome and the rest of the cytoplasm was assumed to be zero. (ii) The DNA Kuhn length was assumed to be 100 nm (i.e., persistence length = 50 nm). This is a popular assumption, but in the context of the cell, it neglects the impact of the high ionic strength and presence of multivalent ions in the cytoplasm. Indeed, an underappreciated result in the aforementioned electron microscopy studies was the observation that expansion of the released chromosomes after cell lysis was considerably reduced with increasing salt concentrations (Kavenoff and Bowen, 1976). Furthermore, the phase diagram in Figure 4B shows that for a Kuhn length of 100 nm (persistence length = 50 nm), the average mesh size would be 13 nm. Such a small average mesh size is incompatible with our findings (Figure 3) or the well-known coupling between transcription and translation in bacteria, which requires ribosomal subunits (~25 nm) to penetrate the nucleoid (see below). (iii) The chromosome was assumed to be linear, neglecting the small relative decrease in polymer size expected after adjusting for its circularity (Casassa, 1965; Kramers, 1946; Rubinstein, 2003; Zimm and Stockmayer, 1949).

After accounting for the solvent quality of the cytoplasm, the estimated rigidity of the DNA under physiological conditions and the circularity of the chromosome (STAR Methods), we found that the theoretical estimation of the chromosome goes down to ~9 μm^3^, which is only ~10-fold (as opposed to 1,000-fold) larger than its experimentally determined size in cells (Figure 8, inset). The remaining compaction needed is likely achieved by other compacting factors such as DNA supercoiling, NAPs, nucleoid-associated RNAs and macromolecular crowding (Cunha et al., 2001; Dame, 2005; de Vries, 2010; Hammel et al., 2016; Jeon et al., 2017; Jun, 2015; Macvanin et al., 2012; Odijk, 1998; Pelletier et al., 2012; Qian et al., 2017; Shendruk et al., 2015; Wegner et al., 2016; Wu et al., 2019; Yang et al., 2020; Yoshikawa et al., 2010; Zhang et al., 2009; Zimmerman, 1993; Zimmerman and Minton, 1993), though some of these factors may also contribute to the poor solvent quality of the cytoplasm (see below).

### Nucleoid mesh size

The membrane-less nucleoid not only stores genetic information but also functions as a mesh-like physical barrier that can affect the diffusion and spatial distribution of cytoplasmic components. Therefore, the mesh size of the nucleoid is an important length scale to consider. By assessing the degree of nucleoid exclusion for fluorescent probes of varying sizes (25-150 nm), we estimate the apparent average nucleoid mesh size to be around 50 nm in exponentially growing *E. coli* cells (Figure 3). This length scale has several physiological implications. For instance, it allows cytoplasmic components of sizes well below 50 nm (e.g., metabolites, proteins, most protein complexes) to diffuse unimpededly in the presence of the nucleoid. As a result, most of the cytoplasm is “well mixed”, facilitating biochemical processes. Since the sizes of free ribosomal subunits are ~23 to 26 nm (Boublik, 1985; Verschoor et al., 1985; Zhu et al., 1997), the 50-nm exclusion size of the nucleoid also enables the coupling between transcription and translation by allowing free ribosomal subunits to reach nascent mRNAs within the nucleoid through diffusion (Sanamrad et al., 2014). If the cytoplasm was an ideal or good solvent, our work suggests that the observed high concentration of the DNA (7 mg/ml) within the nucleoid region would lead to an average mesh size of 20 nm or smaller (Figure 4B) due to the relatively high overlapping of DNA segments with each other (Figure 5A; Video S2). Such a small average mesh size would, at least partly, exclude ribosomes and their subunits from the nucleoid, hindering co-transcriptional translation. Under this scenario, translation would have to primarily take place after transcription has completed and bare (ribosome-free) mRNAs have diffused out of the nucleoid. Such a decoupling between transcription and translation would not only lead to a delay in protein synthesis, but may also shorten the lifetime of mRNAs, as ribosome binding reduces mRNA degradation (Deana and Belasco, 2005; Dreyfus, 2009).

Our data predict that the diffusion and spatial distribution of cytoplasmic components larger than 50 nm are impacted by the presence of the nucleoid in the cytoplasm. We expect that some (possibly most) polyribosome species belong to this category. This reasoning is consistent with the observation that ribosomes, which are mostly found as polyribosomes (Dai et al., 2016; Forchhammer and Lindahl, 1971; Phillips et al., 1969; Varricchio and Monier, 1971), display timescale-dependent sub-diffusive dynamics (Gray et al., 2019), as we may expect for objects that experience caging and uncaging events (Brangwynne et al., 2009; Cai et al., 2011; Guo et al., 2014; Tseng et al. 2004; Wong et al., 2004). Furthermore, other experiments suggest that gene loci and active RNA polymerases (and associated nascent mRNAs) preferentially relocate to the nucleoid periphery during transcription and translation (Cabrera and Jin, 2003; Libby et al., 2012; Stracy et al., 2015; Yang et al., 2019). Once polyribosomes escape the nucleoid, they are less likely to get back inside the nucleoid due to their large size. As a consequence, nucleoid exclusion of polyribosomes leads to ribosome enrichment outside the nucleoid (Azam et al., 2000; Bakshi et al., 2012; Gray et al., 2019; Robinow and Kellenberger, 1994; Sanamrad et al., 2014). The spatial heterogeneity of the DNA density across the nucleoid implies that some regions of the nucleoid are more compacted than others. This is consistent with the spatial heterogeneity of (poly)ribosomes seen by cryo-electron tomography (Figure 6 and Figure S4) and fluorescence microscopy (Figure 7). We expect that the motion and spatial distribution of other large (>50 nm) cellular components, such as storage granules, plasmids, bacterial microcompartments (protein-based metabolic organelles) and stress-induced protein aggregates, are also impacted by the nucleoid, for which there is some evidence (Chowdhury et al., 2014; Henry and Crosson, 2013; Racki et al., 2017; Reyes-Lamothe et al., 2014; Wang et al., 2016; Winkler et al., 2010).

### Chromosome domain organization

The bacterial chromosome is not simply stuffed into the cell; it is instead organized over several length scales (Dame et al., 2020; Verma et al., 2019). Notably, recent Hi-C experiments have shown that the chromosome is partitioned into various domains (CIDs) in which DNA loci preferentially interact with each other (Le et al., 2013; Lioy et al., 2018; Marbouty et al., 2014; Marbouty et al., 2015; Val et al., 2016; Wang et al., 2017; Wang et al., 2015). How these domains form is not fully understood. Our Monte Carlo simulations show that at the physiological DNA concentration of 7 mg/ml, a poor solvent leads to high spatial heterogeneity in DNA density within the nucleoid (Figure 5A-B; Video S2), which is consistent with 3D super-resolution fluorescence microscopy observations of fluorescently-labeled nucleoids in *E. coli* (Le Gall et al., 2016; Marbouty et al., 2015; Spahn et al., 2014; Spahn et al., 2018; Stracy et al., 2015). Such spatial heterogeneity is a consequence of the formation of much less dense DNA regions. By calculating the distance between pairs of DNA segments, we found that the dense regions of the simulated chromosomes correspond to neighboring segments that are spatially clustered together (Figure 5C-D), forming domain-like structures of resemblance to CIDs in Hi-C experiments. In contrast, the simulated chromosomes in an ideal or good solvent were much more homogenous in DNA density (Figure 5B), showing very few, if any, domains of significant size (Figure 5C-D). This argues that the poor solvent quality of the cytoplasm promotes the spontaneous formation of chromosomal domains. There is a precedent for such a solvent-driven structural organization of a biopolymer, as protein folding is highly dependent on the poor solvent condition in vitro (Haran, 2012).

In our simulated chromosomes, domain boundaries were able to form at any chromosomal position (Figure S3A). As a result of this stochasticity, boundaries between domains became increasingly less defined as more distance maps of simulated chromosomes were averaged (Figure 5E and Figure S3B). This is in contrast with Hi-C contact maps in which boundaries of CIDs remain visible despite averaging over billions of cells. Our data therefore suggest that the poor solvent quality of the cytoplasm promotes domain organization of the chromosome, but the specific positions of the domain boundaries seen in Hi-C maps must be attributed to other factors. Indeed, high transcriptional activity is an important determinant of CID boundaries (Le et al., 2013; Le and Laub, 2016; Lioy et al., 2018; Marbouty et al., 2015). In our simulated chromosomes, the domains and their boundaries correspond to DNA regions of high and low densities, respectively. This also appears to be true inside cells, as DNA at the CID boundaries is less condensed (more extensible) than the DNA within CIDs (Le and Laub, 2016). Besides gene expression, other factors, such as NAP binding and membrane attachment, may also contribute to the predominance of specific domain boundaries across cells (Lioy et al., 2018; Marbouty et al., 2014).

Our observation that distinct domain boundaries vanished upon averaging a large number of simulated chromosomes (Figure 5E and Figure S3B) highlights how variability in domain organization may be missed in Hi-C experiments performed on cell populations. At the single-cell level, the structure of the actual chromosome (e.g., size and position of the domains and their boundaries) may considerably differ across cells or change over time, which would be obscured after averaging all identified contacts between DNA loci across a cell population. Consistent with this idea, single-cell Hi-C experiments in eukaryotes have shown that chromosome folding is highly heterogeneous across cells, with the positions of contact enrichments being highly variable (Flyamer et al., 2017; Nagano et al., 2013; Nagano et al., 2017; Ramani et al., 2017; Stevens et al., 2017; Tan et al., 2018; Tan et al., 2019). It would be interesting to perform similar single-cell Hi-C experiments in bacteria. Based on our results, we envision that the solvent quality of the cytoplasm may contribute to some stochasticity in chromosome conformation.

### What may contribute to the poor solvent quality of the cytoplasm?

There are many ways a solution can be a poor solvent for a given polymer. A poor solvent is a solution in which the net interaction between the polymer and the solvent is energetically unfavorable regardless of the nature of such interactions (e.g., electrostatic, hydrophobic, van der Waals, or hydrogen bonding). Due to the complex chemical composition of the bacterial cytoplasm, the observed poor solvent quality of the cytoplasm is unlikely to originate from a single mechanism. A combination of factors is expected to be in action.

Among these factors, NAPs are attractive candidates. Due to their high abundance and specific interactions with the chromosome, the contribution of NAPs to the poor solvent quality of the cytoplasm may be direct or indirect. In vitro, NAPs have been shown to bend, loop, wrap or bridge DNA segments (Dame et al., 2020; Dillon and Dorman, 2010). These molecular interactions may promote local attraction among nearby chromosomal segments in vivo. In addition, we speculate that NAPs, and potentially other DNA-binding proteins (such as transcriptional regulators), may act as an interface that indirectly modulates the chemical interaction between the chromosome and other cellular components of the cytoplasm. Protein binding to the anionic DNA polymer could, for instance, locally alter the electrostatic potential of the chromosome.

There are several lines of evidence suggesting that RNAs may also contribute to the poor solvent quality of the cytoplasm for the DNA. (i) Rifampicin treatment of *E. coli*, which blocks transcription initiation and leads to mRNA depletion through degradation (Bernstein et al., 2002; Chen et al., 2015; Selinger et al., 2003), results in nucleoid expansion (Bakshi et al., 2014; Bakshi et al., 2012; Cabrera et al., 2009; Cabrera and Jin, 2003; Dworsky and Schaechter, 1973; Pettijohn and Hecht, 1974; Sun and Margolin, 2004). Conversely, chloramphenicol treatment, which stabilizes mRNAs (Lopez et al., 1998; Pato et al., 1973; Schneider et al., 1978), is associated with nucleoid compaction (Bakshi et al., 2014; Bakshi et al., 2012; Cabrera et al., 2009; van Helvoort et al., 1996; Zimmerman, 2002). These observations are often interpreted as a consequence of a loss or gain of polyribosome crowding in rifampicin- or chloramphenicol-treated cells, respectively. The idea is that polyribosomes create a depletion force through a volume exclusion effect. However, a mutually non-exclusive alternative is that the chemical nature of these mRNAs, and not just the size of the polyribosomes, plays a role by affecting the quality of the solvent for the chromosome. (ii) The spatial organization of gene expression in *E. coli* is also consistent with RNAs contributing to the poor solvent quality of the cytoplasm. As mentioned above, although transcription can start within the nucleoid, gene loci have been shown to relocate to the nucleoid periphery during transcription (Libby et al., 2012; Yang et al., 2019). This is consistent with the accumulation of active RNA polymerases at the periphery of the nucleoid and regions of low DNA densities (Stracy et al., 2015). This has led to a model in which actively transcribed gene loci with their associated RNA polymerases and nascent mRNAs segregate away from the nucleoid bulk. Importantly, gene relocation occurs independently of concurrent translation, i.e., without the loading of the bulky ribosomes (Yang et al., 2019). These observations support the idea that the interaction between mRNA and DNA is unfavorable (i.e., repulsive). (iii) Repulsive interaction between mRNA and DNA is also consistent with the finding that high levels of long transcripts drive the local establishment of chromosomal domain boundaries (Le et al., 2013; Le and Laub, 2016; Lioy et al., 2018; Marbouty et al., 2015). At the domain boundary, high transcriptional activity leads to an accumulation of nascent mRNAs. Since these mRNAs are still bound to the DNA through the RNA polymerases, they cannot diffuse away. We envision that in order to minimize the contact with these bound mRNAs, nearby DNA strands retract to the flanking sides, reducing their local density at the boundary while forming denser DNA regions (domains) on both sides of the boundary. Such a DNA retraction would be mainly driven by its unfavorable interaction with the mRNA and not by the steric blocking of ribosomes, as the formation of a domain boundary occurs in the absence of translation (Le and Laub, 2016).

Another potential contributing factor to the poor solvent quality of the cytoplasm is its high ionic strength (Alatossava et al., 1985; Cayley et al., 1991; Kuhn and Kellenberger, 1985; Lusk et al., 1968; Moncany and Kellenberger, 1981; Roe et al., 1998; Schultz et al., 1962). In vitro studies have shown that the net interaction between DNA fragments becomes more attractive with increasing salt concentrations (Nicolai and Mandel, 1989). Divalent Mg^2+^, but not monovalent Na^+^, has been found to not only screen the charges on DNA but also induce attraction among the strands (Qiu et al., 2007). Overall, the poor solvent quality is likely to originate from interspersed and superimposed molecular interactions associated with the complex chemical nature of the bacterial cytoplasm.

## Acknowledgements

We are grateful to Dr. David Baker for the nanocage-producing strain and to Drs. Corey O’Hern, Thierry Emonet and Eric Dufresne for valuable feedback. We also thank the members of the Jacobs-Wagner laboratory for fruitful discussions and for critical reading of the manuscript. This work was partially supported by the National Institute of Allergy and Infectious Diseases (R01AI087946 and R01AI132818 to J.L.). C.J.-W. is an investigator of the Howard Hughes Medical Institute.

## Author Contributions

Conceptualization, Y.X. and C.J.-W.; Methodology, Y.X., I.V.S., Y.C., S.K.G., B.R.P. and C.J.-W. Software, Y.X.; Formal Analysis, Y.X., I.V.S., Y.C. and B.R.P.; Investigation, Y.X., I.V.S., Y.C., S.K.G. and B.R.P.; Data Curation, Y.X., I.V.S., Y.C., S.K.G. and B.R.P.; Writing - Original Draft, Y.X. and C.J.-W.; Writing - Review & Editing, Y.X., I.V.S., Y.C., S.K.G., B.R.P., J.L. and C.J.-W; Visualization, Y.X., I.V.S., Y.C. and C.J.-W.; Supervision, J.L and C.J.-W.; Project Administration, C.J.-W.; Funding Acquisition, C.J.-W.

## Supplemental figure legends

**Figure S1.**
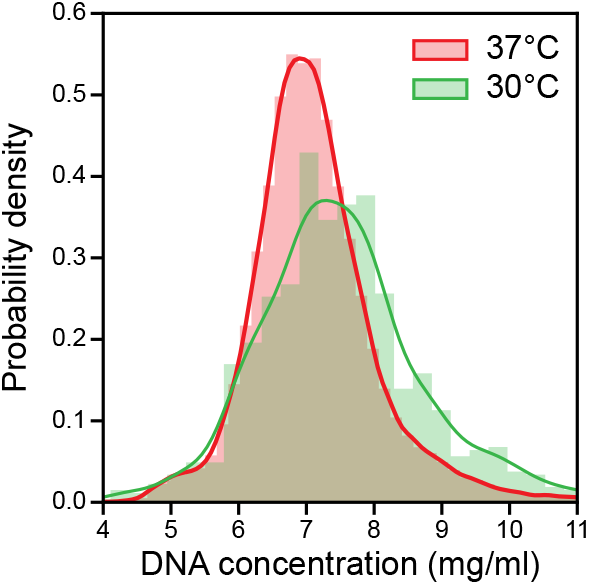
Related to Figure 2. Comparison of DNA concentration measurements at 30°C and 37°C growth temperatures. Probability density functions of the average DNA concentrations estimated within the nucleoid region in B-period cells (CJW6324) grown at either 30°C (green) or 37°C (red).

**Figure S2.**
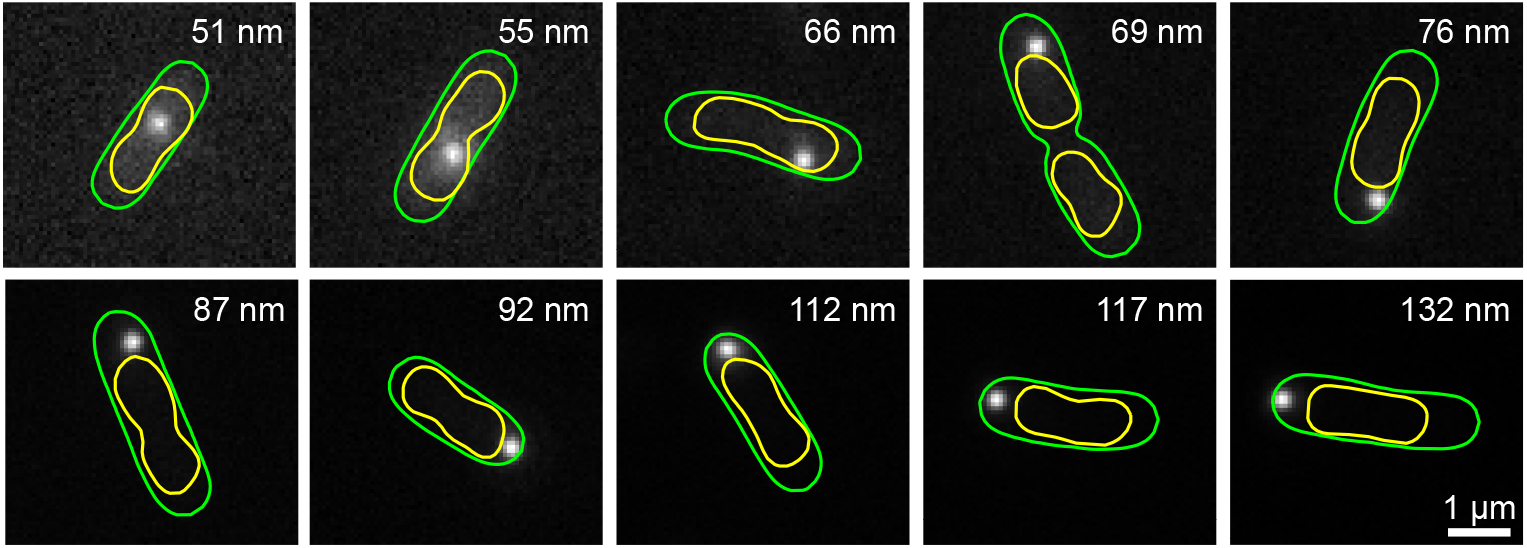
Related to Figure 3. Example GFP-μNS particles of various sizes. Each panel shows a GFP fluorescence image of an *E*. coli cell (CJW4617). The GFP-μNS particles appeared as distinct bright spots. The particle size is indicated in the upper right corner. The green and yellow outlines are the cell and nucleoid contours detected by the Oufti software package.

**Figure S3.**
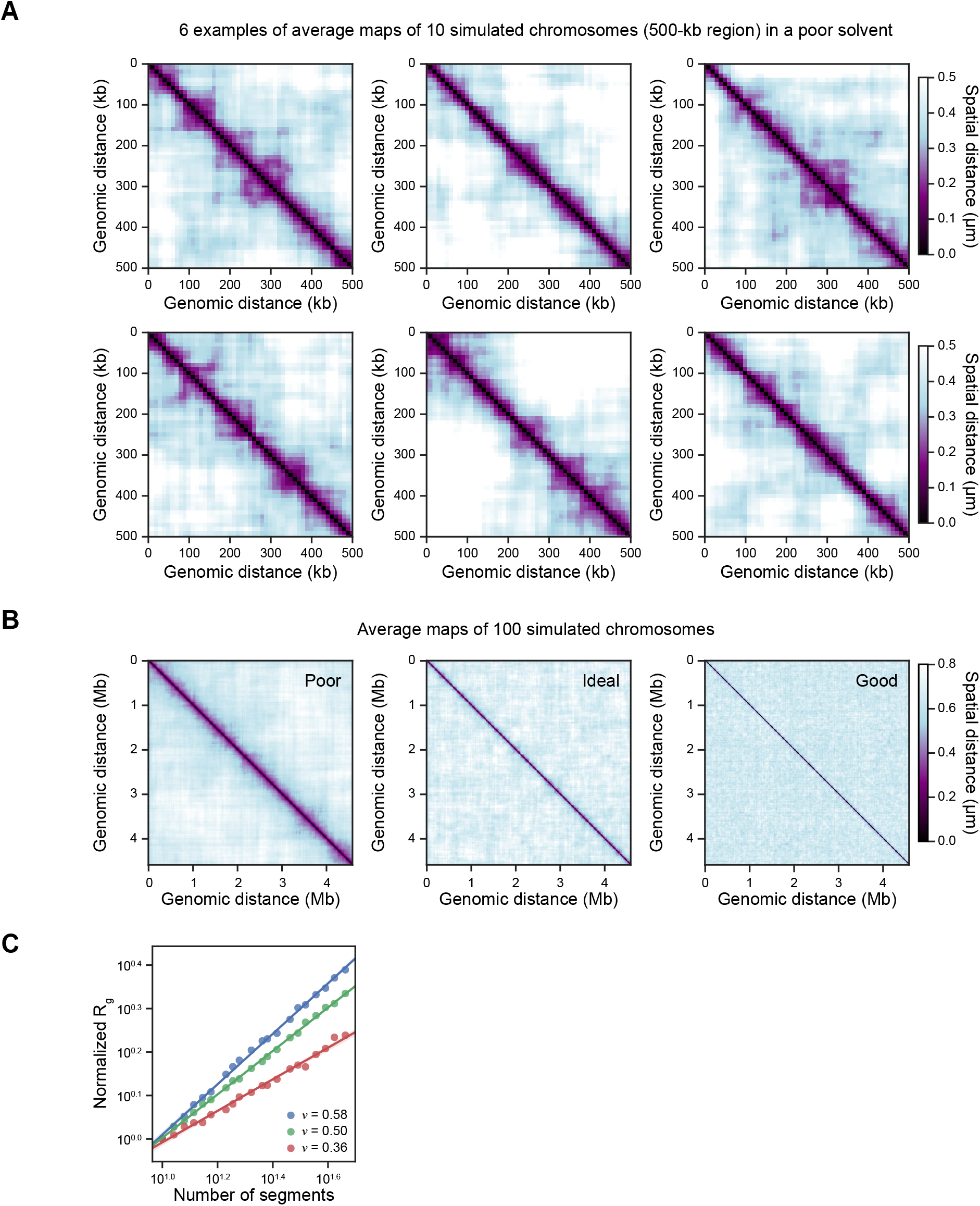
Related to Figure 5. Monte Carlo simulations of *E. coli* chromosomes. A. Six examples of distance maps indicating the spatial separation between any two loci along simulated chromosomes in a poor solvent (*v* = 0.36). Each map is the result of averaging 10 randomly selected simulated chromosomes. B. Distance maps obtained by averaging 100 simulated chromosomes in a poor (*v* = 0.36), ideal (*v* = 0.50) and good solvent (*v* = 0.58). Same as Figure 5E, except for showing the distance maps for the full simulated chromosomes. C. Scaling between the number of segments and the normalized *R_g_* of simulated chromosomes in different types of solvent. For a better comparison in the scaling, the *R_g_* of the simulated chromosomes of different numbers of segments were normalized by the *R_g_* of the chromosome with 10 segments. Each point is based on the average of 200 simulated chromosomes. Note that the scaling was only shown for small length scales because the excluded volume interactions are screened for polymers at large length scales. As a result, the scaling exponent becomes 0.5, regardless of the solvent quality (Rubinstein, 2003).

**Figure S4.**
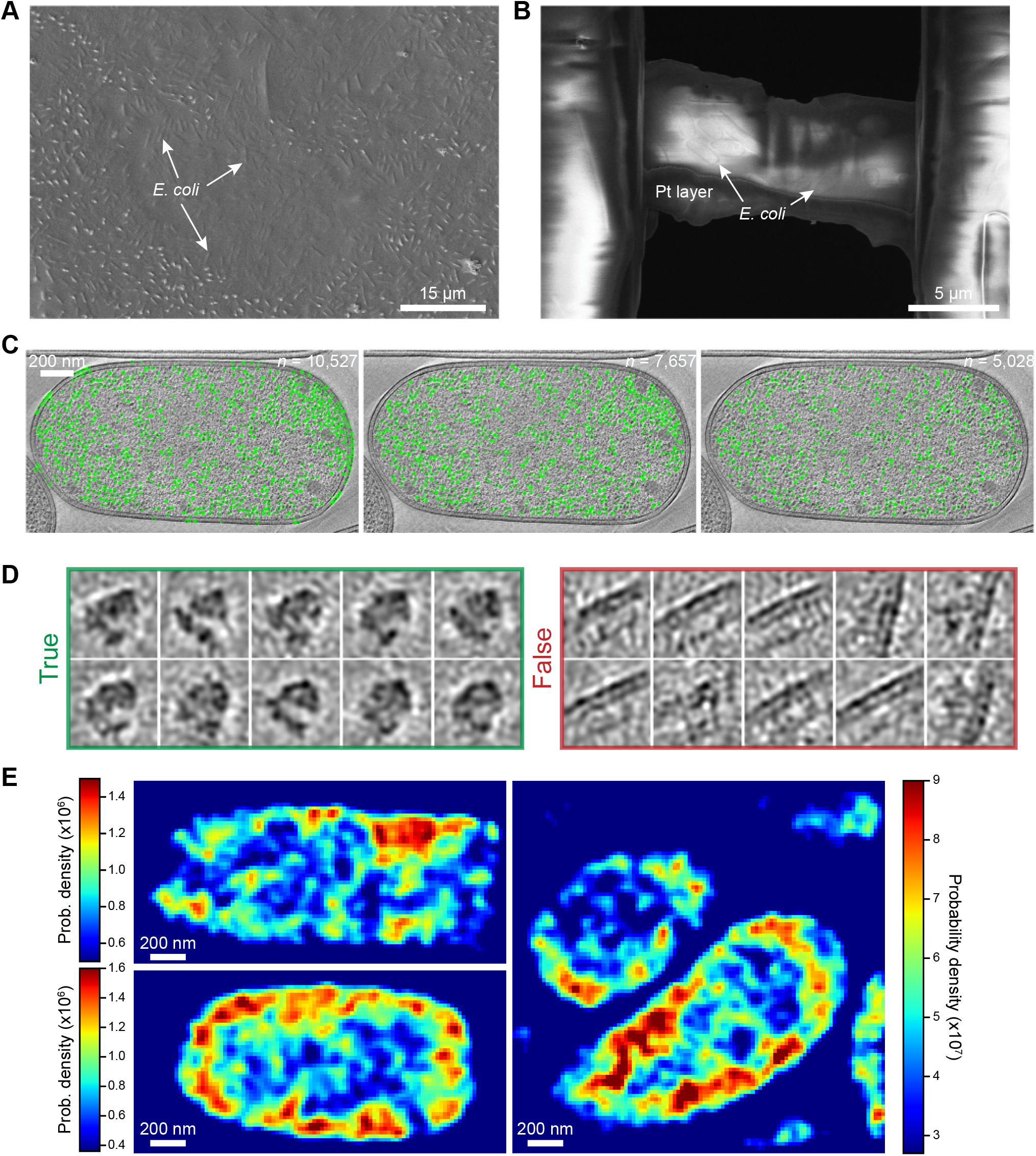
Related to Figure 6. Localization of ribosomes using cryo-ET. A. Representative scanning electron microscope (SEM) image showing the *E. coli* cells on the grid before the cryo-FIB milling. B. Top view of the SEM image showing a lamella after the final polishing in cryo-FIB milling. C. From left to right, each panel represents the same tomogram section overlaid with the ribosomes identified by the initial 3D template search, after the removal of structures in the false class, and after the removal of duplicate structures. In each case, the number of localized ribosome structures (*n*) is indicated in the top right, and their positions are marked by the green circles. D. Representative structures in the true and false classes during classification of the candidate 70S ribosome structures found in the 3D template search. E. Additional examples of probability density maps of ribosome structures localized in tomograms. All maps were Gaussian smoothed (*σ* = 1.2 pixel, 21 nm) for easier visualization of the ribosome regions of low and high densities.

## STAR Methods

### Strains and growth conditions

This study used the following published strains. MG1655 (F-lambda-*ilvG-rfb-50 rph-1*) was used as a wild-type *E. coli* strain (Jensen, 1993). Strain CJW6324 (MG1655 *seqA::seqA-mcherry ftsZ::ftsZ-venus^SW^*) was used to separate the cell population into three distinct groups that corresponded to the cells in the B, C and D periods (Gray et al., 2019). Strain CJW6340 (BL21 Star (DE3) pET29b+ I3-01-sfgfp) was obtained as a kind gift from David Baker’s laboratory at the University of Washington (Hsia et al., 2016). In this strain, the artificially designed protein I3-01 fused to superfolder GFP (I3-01(ctGFP)) was produced from the pET29b+ plasmid in BL21 Star (DE3) *E. coli* cells. I3-01(ctGFP) assembles into dodecahedron nanocage particles (Hsia et al., 2016). Strain CJW4617 (MG1655 Δ*lacZYA::gfp-μNS*) produces GFP-μNS particles of tunable sizes in response to IPTG induction (Parry et al., 2014). Strain CJW7020 (MG1655 *rplA::rplA-msfgfp*) was used to investigate the pixel intensity correlation between DNA and ribosome fluorescence signals (Gray et al., 2019).

To obtain steady-state growth conditions, a sample of an overnight liquid cell culture in stationary phase was diluted by a factor of at least 10,000 in the corresponding fresh growth medium. The culture was then allowed to grow in an incubator shaker (New Brunswick Innova 44), at the indicated temperature, to reach an optical density at 600 nm (OD_600_) of 0.1-0.3 prior to microscopy.

### Light microscopy and image analysis

Unless otherwise specified, all exponentially growing cells were imaged on 1% agarose pads supplemented with growth medium. When appropriate, cultures were incubated with 1 μg/ml 4’,6-diamidino-2-phenylindole (DAPI) for 15 min in growth medium prior to imaging on agarose pads. Phase contrast and epifluorescence imaging were performed on a Nikon Eclipse Ti-E microscope equipped with a phase contrast Nikon CFI Plan Apo DM Lambda 100x oil objective (NA = 1.45), and a SOLA light engine (Lumencor). A Ph3 phase contrast ring was used. The following Chroma filters were used for epifluorescence imaging: DAPI (excitation D390/22x, dichroic T425lpxr, emission ET460/50m), GFP (excitation ET470/40x, dichroic T495lpxr, emission ET525/50m), mCherry/TexasRed (excitation ET560/40x, dichroic T585lp, emission ET630/75m). The microscope was controlled by the NIS-Elements AR software.

Using the open source software Oufti (Paintdakhi et al., 2016), cell and nucleoid outlines were constructed based on the phase contrast and epifluorescence images, respectively. When appropriate, cells were classified into the B, C and D cell-cycle periods based on two cell features: SeqA-mCherry pattern and cell area, as described before (Gray et al., 2019).

### Derivation of the nucleoid mesh size

The correlation length *ξ* (i.e., mesh size) of a semidilute polymer solution is defined as *ξ* = *R_g_*(*c*\*/*c*)^*v*/(3*v*−1)^ (Cooper et al., 1991; Dasgupta et al., 2002; Gennes, 1979; Rubinstein, 2003), where *R_g_* is the radius of gyration, *c* is the polymer concentration, and *v* is the Flory exponent. The overlap concentration *c*\*, above which polymer segments overlap to form a mesh can be written as 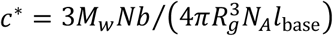 (Mutch et al., 2007; Ramakrishnan et al., 2002), where *M_w_* is the average molecular weight of the base pair (650 g/mol) (Lee et al., 2012; Peale et al., 1989; Ratilainen et al., 2001), *N* is the number of polymer segments, *b* is the Kuhn length, *l*_base_ is the size of a base pair (0.34 nm) (Diekmann et al., 1982; Yonemura and Maeda, 1982), and *N_A_* is the Avogadro’s number. For a circular polymer, 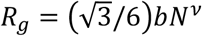 (Rubinstein, 2003). Plugging both the *R_g_* and *c*\* into the above definition, we have 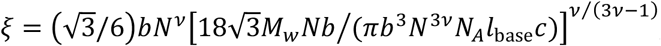, which can be simplified to yield Eq. 1 above: 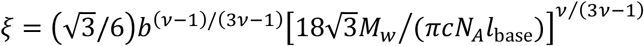. To verify the dimension of this result, we note the term 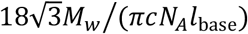 has a dimension of area. Since *b* has a dimension of length and (*v* – 1)/(3*v* – 1) + 2*v*/(3*v* – 1) = 1, *ξ* correctly has a dimension of length.

### Estimation of the average DNA concentration within nucleoids of *E. coli* in the B period

*E. coli* strain CJW6324 was grown at 37°C in M9 minimal medium supplemented with glycerol (0.2%). To estimate the average DNA concentration within the nucleoid regions, B-period cells (with a single chromosome) were first identified (see above). Given that the average molecular weight of a nucleotide base pair is 650 g/mol (Lee et al., 2012; Peale et al., 1989; Ratilainen et al., 2001) and the *E. coli* genome size is about 4.6 million base pairs (Blattner et al., 1997), the total DNA mass in a single chromosome is around 5 femtograms. The nucleoid volume was estimated as *V*_nucleoid_ = *V*_cell_(*A*_nucleoid_/*A*_cell_)^3/2^, where *V*_nucleoid_, *V*_cell_, *A*_nucleoid_, *A*_cell_ are the volumes and areas of the nucleoid and cell. The cell volume was calculated by accumulating the volume of slices defined by neighboring pairs of points along the cell outlines (Paintdakhi et al., 2016) (see also documentation on oufti.org). The nucleoid and cell areas were calculated by summing up the areas of the polygons defined by the nucleoid and cell outlines. The power 3/2 was used to convert the estimated nucleoid area fraction into a volume fraction. The nucleoid volume estimation was performed at the single-cell level, as was the DNA concentration calculation, which was simply a division between the estimated DNA mass and the nucleoid volume.

### Estimation of the average DNA concentration within nucleoids of *E. coli* in nutrient-rich conditions

Bulk studies using the diphenylamine colorimetric assay and cell counting reported that the average DNA mass per cell under nutrient-rich conditions, glucose, glucose plus casamino acids and rich defined medium plus glucose as carbon sources, were around 10, 14 and 30 fg, respectively (Basan et al., 2015). The average cell sizes among these growth conditions (i.e. 2.32, 3.33, 6.60 μm^3^, respectively) were also shown to scale linearly with the average DNA mass per cell (Basan et al., 2015). Taking into consideration of the strong scaling between the nucleoid size and cell size (Gray et al., 2019), we estimated the average DNA concentration within the nucleoid to be 7.3, 7.1 and 7.7 mg/ml under these three nutrient-rich growth conditions. Therefore, when taken together, the DNA concentration within the nucleoid among these nutrient-rich conditions is 7.4 ± 0.2 mg/ml.

### Single-particle tracking experiments

GFP-tagged nanocage particles were produced through basal expression of the GFP-tagged protein subunit I3-01 in *E. coli* cells (CJW6340) grown at 30°C in M9 minimal medium supplemented with glycerol (0.2%), casamino acids (0.1%) and thiamine (1 μg/ml). Phase contrast images were taken first to define cell outlines. Time-lapse fluorescence images were then acquired at a frame rate of 100 ms. Nanocage particles were localized and tracked using the MATLAB class SPT (see supplementary code). Briefly, an image was first smoothed by a band-pass filter. Potential regions containing the particles were then segmented based on an intensity threshold. Based on the region sizes, background noise pixels or particle clusters were filtered out. Within each of the remaining segmented regions, the brightest pixel was located. On the brightest pixel and its eight surrounding pixels, the pixel intensities were fitted by a 2D Gaussian distribution, whose center was taken as the estimated particle position. The particle locations were linked into trajectories using an algorithm described previously (Crocker and Grier, 1996). The 2D mean squared displacements (MSD) plot were calculated as MSD(*τ*) = 〈(*r*(*t* + *τ*) – *r*(*t*))^2^〉, where *τ* is a given time delay and *r*(*t*) and *r*(*t* + *τ*) denote the particle positions before and after the given time delay. Similarly, GFP-μNS particles were produced through IPTG induction (50-200 μM) for a duration of 30-120 min, in *E. coli* cells (CJW4617) grown at 30°C in M9 minimal medium supplemented with glycerol (0.2%), casamino acids (0.1%) and thiamine (1 μg/ml). GFP-μNS particles were detected using SpotFinder (Sliusarenko et al., 2011). Particle positions of both nanocage-GFP and GFP-μNS particles relative to the cell coordinates were calculated using the function projectToMesh in Oufti and normalized by cell width and length. The relative particle positions were used to generate histograms to demonstrate the probability density of relative particle locations inside the cell.

### Estimation of GFP-μNS particle sizes

The size estimation of the GFP-μNS particles was performed as previously described (Parry et al., 2014). This method establishes a relation between the fluorescence intensity (*I*) of GFP-μNS particles and their absolute sizes (*d*). The particle fluorescence intensity (*I*) was calculated by integrating the volume below the fitting bivariate normal distribution that was used to determine the particle location: *I* = 2*πAσ_x_σ_y_*, where *A* is the amplitude of the fitting distribution, and *σ_x_, σ_y_* are the standard deviations of the distribution in X and Y dimensions, respectively. Assuming the total fluorescence intensity of a particle is proportional to its volume, and the volume is a cubic function of its size (e.g., for a sphere), we further calculated the cubic root of the fluorescence intensity: 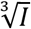.

The absolute sizes of GFP-μNS particles were estimated as previously described (Parry et al., 2014). Briefly, cells expressing GFP-μNS were lysed with T7 phages, resulting in the release of the GFP-μNS particles in solution. Cell debris were removed by centrifugation at 4,000 g. Released GFP-μNS particles were then mixed with fluorescent microspheres of known nominal size (110 nm, FluoSpheres F8803, Invitrogen), and their diffusion coefficients were determined using single particle tracking microscopy. We defined the fluorescence-derived relative particle size 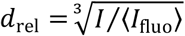 for the GFP-μNS particles, where 〈*I*_fluo_〉 denotes the average fluorescence intensity of the fluorescent microspheres. Based on the Stokes-Einstein equation, the relative particle size and the corresponding diffusion coefficient is inversely proportional: *D* = *kT*/(3*πηd*_rel_), where *k* is the Boltzmann constant, *T* is the absolute temperature, and *η* is the viscosity of the solution. The absolute sizes of the GFP-μNS particles were estimated as *d* = *d*_fluo_ *D*_fluo_ *α*/*D*, where *d*_fluo_ = 110 nm and *α* = *d*/*d*_rel_ is a calibration constant, which was found to be 685 nm.

### Monte Carlo simulation of chromosome conformation

The entire *E. coli* chromosome (4.6 million base pairs, contour length = 1564 μm) was modeled as a chain of 26,066 segments, each of which has a length of 60 nm (to reflect the assumed Kuhn length). To achieve the experimentally determined average DNA concentration within the nucleoid region, we confined the entire chromosome within a spherocylindrical space with a volume (0.70 μm^3^) equal to the experimentally observed average nucleoid volume in B-period cells. In an ideal solvent where the net interaction between the polymer and the solvent is neutral, there is no preference for a segment on the polymer to interact more or less with other segments. In the simulation, at the end of each step, a next-step segment was uniformly sampled from a spherical surface, the center and radius of which are the end of the current step and the assumed Kuhn length, respectively. For a polymer in a poor or good solvent, the segments on the polymer chain ‘prefer’ interacting among themselves (poor solvent) or with the solvent (good solvent). To model the inter-segmental attraction and repulsion, instead of sampling the next-step segments uniformly anywhere on the spherical surface, restrictions were applied on the range of inclination and azimuth of the spherical surface from which next-step segments (i.e., angles between the consecutive segments) could be sampled. By verifying the power-law scaling between the number of segments and the radius of gyration of the simulated chromosomes, the inter-segmental angles were constrained within empirically determined ranges (Figure S3C). Specifically, we found that the Flory exponents (0.36 and 0.58) could be achieved when the inter-segmental angles were constrained to be greater than 11π/18 and smaller than π/2, respectively.

Similarly, in order to compare the degree of chromosome compaction as a result of different types of solvent, the *E. coli* chromosome was simulated in free space 1,000 times for each type of the solvent. The radius of gyration of each simulated chromosome was calculated as 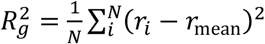, where *R_g_* is the radius of gyration, *r_i_* is position of *i*-th segment, *r*_mean_ is the average position of all segments and *N* is the total number of segments. To account for the circular geometry of the *E. coli* chromosome, the resulted *R_g_* was divided by a factor of 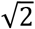 to get 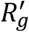. The volume of the simulated nucleoid in each type of solvent was calculated as 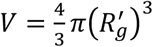.

### Construction of distance maps for simulated DNA segments

The simulated chromosomes were binned by approximately 10 kb (56 DNA segments), and the average positions of the segments within each bin were calculated. The binning was applied to reduce the computational complexity of calculating the pair distance between DNA segments by more than three orders of magnitude. A matrix of pair distances, where the *i*-th row and *j*-th column represent the Euclidean distance between the *i*-th and *j*-th bins, was constructed for every simulated chromosome. A chosen number (10 or 100) of such matrices were averaged together to generate distance maps representing the average pair distances between the binned segments, which were then visualized as images in which the individual pixels correspond to the distances.

### Preparation of the cryo-EM grids

*E. coli* wild type cells (MG1655) were grown at 37°C in M9 minimal medium supplemented with glucose (0.2%), casamino acids (1%) and thiamine (1 μg/ml). Nutrient-rich growth conditions were chosen over nutrient-poor condition to increase the intracellular concentration of ribosomes (Dai et al., 2016; Forchhammer and Lindahl, 1971; Phillips et al., 1969; Varricchio and Monier, 1971) thereby decreasing the likelihood that any observed heterogeneity in ribosome distribution in the cell is due to random clustering. Cells were then harvested and concentrated between 30- and 150-fold by centrifugation (6,000 g for 5 min). The concentrated culture sample was deposited onto freshly glow-discharged holey carbon grids (Quantifoil). The grids were then blotted with the filter paper for ~3-5 s before getting plunge-frozen into liquid ethane, using a custom gravity-driven plunger apparatus described previously (Liu et al., 2009; Zhao et al., 2013).

### Cryo-FIB milling

The plunge-frozen grids were clipped into cryo-FIB AutoGrids and mounted into the specimen shuttle under liquid nitrogen. An Aquilos cryo-FIB system (Thermo Fisher Scientific) was used to mill the samples to produce thin lamellae. The samples were first sputter-coated with Pt (1 kV, 15 mA, 15 s) to improve the overall sample conductivity, and then were deposited by an organometallic Pt layer (4-5 μm thick) using the gas injection system for the sample protection. Lamellae were produced using the gallium ion beam at 30 kV with stage tilt angle around 17°. The ion beam current was reduced according to the lamella thickness (t) during the milling process: 0.5 nA for t ≥ 3 μm, 0.3 nA for t ≥ 1 μm, 0.1 nA for t ≥ 700 nm, 0.05/0.03 nA for final polishing. Afterwards, a thin Pt layer was sputter-coated (1 kV, 10 mA, 5 s) on the lamella to prevent possible charging issue during the cryo-ET imaging.

### Cryo-ET data acquisition and tomogram reconstruction

The cryo-FIB lamellae were transferred to a 300 kV Titan Krios electron microscope (Thermo Fisher Scientific) equipped with a Direct Electron Detector and energy filter (Gatan). The program SerialEM (Mastronarde, 2005) was used to collect single-axis tilt series around −6 μm defocus, with a cumulative dose of ~100 e-/Å covering angles from −60° to 60° (2° tilt step). Images were acquired at 26,000’ magnification with an effective pixel size of 5.457 Å at the specimen level. All recorded images were first drift corrected by the software MotionCor2 (Zheng et al., 2017) and then stacked by the software package IMOD (Kremer et al., 1996). All tilt series were then aligned by IMOD with the patching tracking method. The program Gctf (Zhang, 2016) was used to determine the defocus of each tilt image in the aligned stacks, and the function “ctfphaseflip” in IMOD was used to perform the contrast transfer function (CTF) correction for the tilt images. Tomograms were then reconstructed in IMOD using the CTF-corrected and aligned stacks.

### Ribosome localization in the cryo-ET tomograms

The software package emClarity (Himes and Zhang, 2018) was used to perform the 3D template search to identify ribosomes from the tomograms. A low-pass filtered (~ 4-nm resolution) *E. coli* 70S ribosome structure (Fu et al., 2019) was used as a reference for the template search (Figure S4C). Initially, up to 15,000-20,000 sub-volumes were initially selected from each tomogram. Subsequently, the 3D classification and alignment were applied to all sub-volumes by the software package i3 (Winkler, 2007; Winkler et al., 2009) with no binning. Sub-volumes of the particles in the false class were removed (Figure S4D). After that, for every pair of localized structures, the separation distance between their center mass was calculated. Two structures in a pair were considered duplicates, if the distance between their centers of mass was smaller than 20 nm, and the structure that had a lower cross-correlation score with the global average structure was removed. This process repeated itself until no duplicate structure was found. The program UCSF ChimeraX (Goddard et al., 2018) was used for the 3D surface rendering of subtomogram average structure of ribosome. The accuracy of the localization of ribosomes was verified by both visual inspection and comparison of the average structure obtained from the localized ribosomes to the reference structure.

### Pixel intensity correlation between the DNA and ribosome fluorescence signals

*E. coli* cells (CJW7020) were grown at 37°C in M9 minimal medium supplemented with glucose (0.2%), casamino acids (1%) and thiamine (1 μg/ml). The nucleoid was visualized through DAPI straining. Fluorescence signals from DAPI and L1-msfGFP were acquired. The nucleoid outlines were constructed using objectDetection in the software package Oufti (Paintdakhi et al., 2016). In order to exclusively quantify the pixel intensity correlation *within* the nucleoid region, the nucleoid outlines were shrunk by 2 pixels from the original boundaries using the built-in MATLAB function polybuffer. The shrunk nucleoid outlines enclosed, on average, about 30% of the inner most pixels defined by the original outlines. Image masks were constructed using the shrunk nucleoid outlines, and pixels from both the DAPI and GFP images were extracted. Correlation scores (Spearman’s *ρ*) were then calculated between the intensities of DAPI and GFP pixels using the built-in MATLAB function corr.

## Supplemental video legends

**Video S1. Related to Figure 3. Diffusion of a 25-nm nanocage-GFP particle**

This video shows an example trajectory of nanocage-GFP particle diffusing in the cytoplasm of an *E. coli* cell (CJW6340). The left panel shows the original GFP fluorescence signal from the particle, and the right panel shows the additional annotation based on the single-particle tracking results: the particle center was denoted by the red point, and the yellow tail represents a short trajectory indicating the particle positions within the last 5 frames.

**Video S2. Related to Figure 5. Simulated chromosomes in different types of solvent.**

This video shows a 3D overview of chromosomes simulated in confinement of a spherocylinder with a volume equal to that of an average nucleoid (~0.7 μm^3^). From left to right, the solvent was assumed to be poor (*v* = 0.36), ideal (*v* = 0.50) and good (*v* = 0.58). In all three cases, the Kuhn length of the DNA was assumed to be 60 nm.

**Video S3. Related to Figure 6. Tomogram of the *E. coli* lamella**

This video shows the tomogram of an *E. coli* lamella obtained after the FIB milling. The lamella had an average thickness of 155 nm, and 5,028 ribosomes were detected in it. Based on their corresponding positions and orientations identified in the analysis, the ribosome structures (colored yellow) were placed back into the tomogram for visual inspection. The video shows individual tomogram sections along the Z-axis.

**Video S4. Related to Figure 6. Comparison between the reference and the average structure of localized 70S ribosomes**

This video shows a 3D comparison between the low-pass filtered (~3 nm resolution) reference structure (grey) obtained from (Fu et al., 2019) and the average structure (yellow) based on all ribosomes localized in the tomogram shown in Figure 6B and Video S3.

**Video S5. Related to Figure 8. Simulated chromosomes in free space in different types of solvent**

This video shows a 3D overview of chromosomes simulated in free (unbound) space in different types of solvent. From top to bottom, the solvent was assumed to be poor (*v* = 0.36), ideal (*v* = 0.50) and good (*v* = 0.58). In all three cases, the Kuhn length of the DNA was assumed to be 60 nm

## Analysis and simulation code availability

All code used for analysis in this study can be found at https://github.com/lacobsWagnerLab.

